# PETRO enables robust estimation of on-target perturbation effects in Perturb-seq

**DOI:** 10.64898/2026.10.15.759256

**Authors:** Zhongxuan Sun, Hyunseung Kang, Sündüz Keleş

**Affiliations:** Department of Biostatistics and Medical Informatics, University of Wisconsin–Madison; Department of Statistics, University of Wisconsin–Madison

## Abstract

Perturb-seq uses multiple guides per target as replicates, but off-target effects can bias target-level estimates even with efficient target perturbation. We present PETRO, which recovers the on-target perturbation effect while allowing, for each outcome, an unknown minority of guides to have arbitrary off-target effects. Across CRISPRa and CRISPRi datasets, sequence-associated off-target bias persisted after pooling cells across guides and nearly doubled in one CRISPRa dataset after a published guide filter. PETRO largely removed this bias and reduced transmission of effects from independently flagged promiscuous guides. In simulations, PETRO nearly matched pooling in power without off-target effects; with off-target effects, pooling remained biased as cell numbers increased, whereas PETRO recovered the on-target perturbation effect. In cortical Perturb-seq, most ZNF219 pathway enrichments from pooling rested on one guide and did not survive majority-supported aggregation, unlike those for targets with concordant guides. PETRO enables robust target-level Perturb-seq analysis from replicate-guide agreement.

## Main

Perturb-seq couples pooled CRISPR perturbations with single-cell RNA sequencing to profile transcriptome-wide responses at scale, typically using several guide RNAs per intended target [1–3]. Methods such as DESeq2, SCEPTRE, Mixscale and causarray estimate and test expression changes associated with genetic perturbations [4–7]. When information from several guides assigned to the same target is combined, these analyses are often interpreted as estimating a target-level perturbation response. That interpretation becomes ambiguous when individual guides produce additional effects not attributable to the intended perturbation.

Recent studies show that a guide can successfully perturb its intended target while also altering expression through off-target binding [3, 8–10]. A guide’s effect can therefore contain both an on-target component, induced by perturbing the intended target, and an off-target component, induced by unexpected off-target activity. Target-level estimates are meant to capture only the on-target component, but accurate estimation of guide effects does not disentangle the two components. Off-target activity can thus distort target-level estimates, and the biological conclusions drawn from them, even when the intended target is efficiently perturbed.

Multiple guides provide replication for addressing this attribution problem. The most common approach pools cells carrying guides for the same target or averages the estimated guide effects [1, 5, 11]. Such aggregation dilutes each guide’s off-target effect according to its weight but eliminates bias in the target-level estimate only if the weighted effects cancel. More cells therefore narrow uncertainty around a biased estimate rather than recovering the on-target component. Alternatively, sequence or perturbation-profile evidence can identify suspicious guides for removal [3, 9, 10]. These diagnostics are valuable for assessing off-target liability, but filtering flagged guides does not ensure unbiased target-level estimates: retained guides need not be valid for every outcome, and renormalizing their weights can increase the remaining bias.

Model-based approaches provide another route. Canon formulates guide assignments as instrumental variables and estimates the causal effect of target-gene expression on an outcome (the regulatory effect) while allowing additional off-target effects [12]. This estimand differs from the on-target perturbation effect that conventional guide pooling aims to estimate. Canon separates on-target and off-target effects through a spike-and-slab prior in which nonzero off-target effects are mean-zero Gaussian draws independent of guide strength. Here we ask a different question arising directly in standard multi-guide Perturb-seq analysis: when a minority of replicate guides may carry arbitrary outcome-specific deviations, what on-target perturbation effect is identified by guide agreement?

We present PETRO (perturbation effect estimation robust to off-target bias), a framework for estimating the on-target perturbation effect under a majority rule. For each target-outcome pair, the primary model assumes that all active guides share the same on-target perturbation effect, while invalid guides additionally carry off-target effects. If a strict majority of guides are valid, the on-target perturbation effect equals the median of the guide effects. It is therefore identified without knowing which guides are valid, and deviations in the invalid minority may be arbitrary in sign and magnitude [13, 14]. Instead of removing flagged guides for all outcomes, PETRO estimates this shared effect by robustly combining guide-level estimates with outcome-specific weights reflecting precision and agreement. Because it takes guide-level effects and their uncertainty as input, the upstream outcome model can be chosen to suit the data and design. A proportional extension estimates the regulatory effect, enabling direct comparison with Canon. Held-out-guide prediction favored neither model consistently (Supplementary Fig. S1), so PETRO retains both but prioritizes the common-effect model to match the shared-effect interpretation of conventional pooling [15].

Across published CRISPRa and CRISPRi screens, sequence-associated bias persisted after conventional pooling, and post hoc guide filtering increased residual sequence-associated structure among retained guides. PETRO attenuated this bias without using sequence annotations and reduced transmission of effects from independently flagged promiscuous guides. In simulations without off-target effects, PETRO nearly matched pooling in power. With one off-target guide, pooling stayed biased as cell numbers increased, whereas PETRO’s bias approached zero. Gaussian approximations can inflate type I error in sparse Perturb-seq data [5]. PETRO supports both Gaussian-based and permutation inference; on null data built from control cells, permutation tests remained calibrated whereas Gaussian-based tests did not. Finally, in a cortical Perturb-seq study [11], pooling masked conflicting lineage and transcriptional effects among ZNF219 guides with comparable knockdown. PETRO retained few pooled ZNF219 pathway enrichments but most for concordant ARX and NR2E1, the study’s other two top candidates.

## Results

### Overview of PETRO

PETRO is a causal inference framework for Perturb-seq that uses multiple guides per target as a source of identification in the presence of off-target effects. Under the majority rule, which assumes that more than half of a target’s guides are valid for each outcome, the on-target perturbation effect can be identified without knowing which guides are valid, while the off-target perturbation effects of the invalid minority are unrestricted. For each target-outcome pair, PETRO combines estimated guide effects and their covariance to return a target-level effect estimate, parametric or permutation-based *P* value, and guide-specific weights reflecting estimation precision and agreement (Fig. 1a and Methods).

**Fig. 1.**
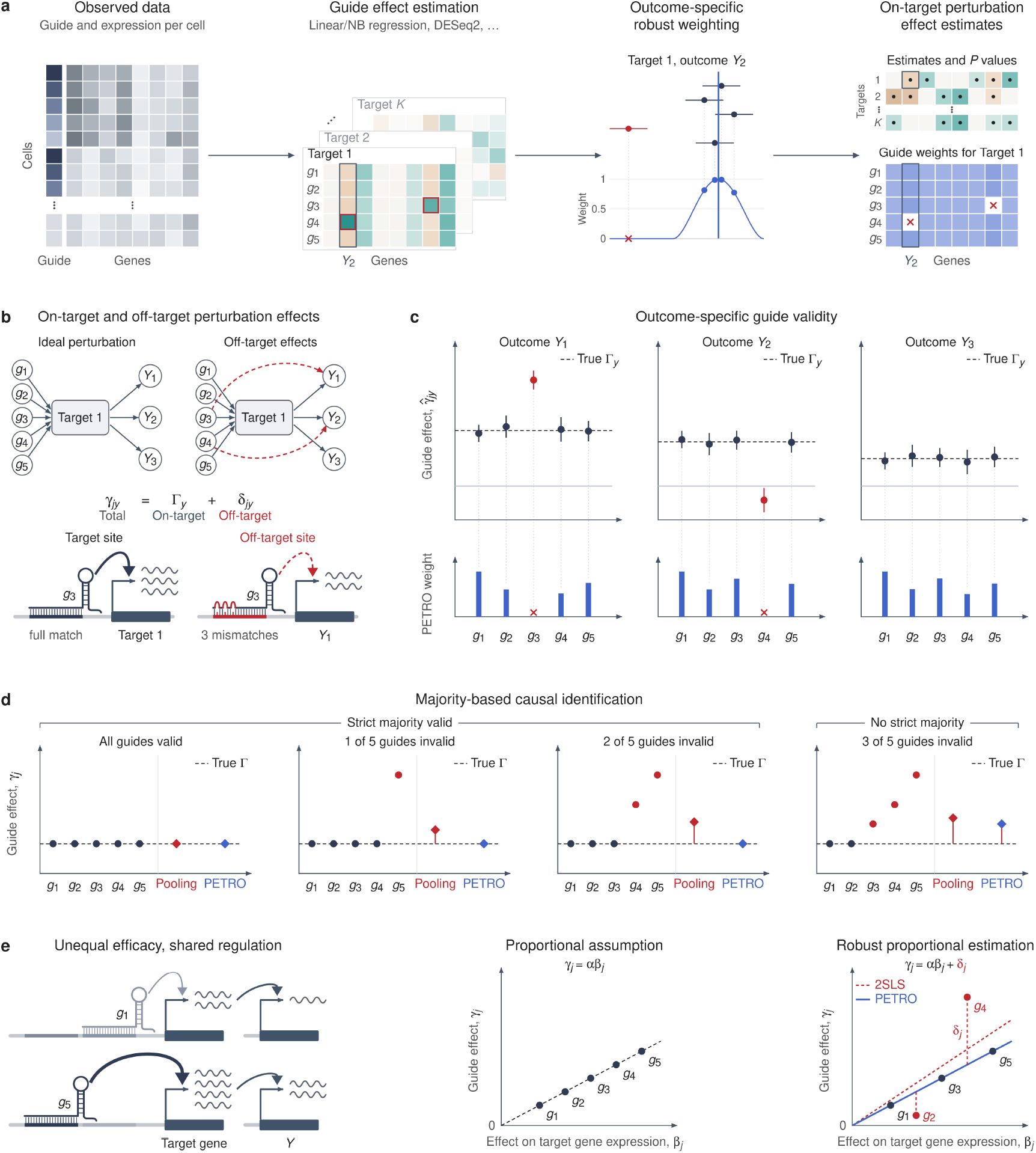
PETRO estimates on-target perturbation effects under the majority rule. **a**, Workflow. Guide effects on each outcome relative to control cells, with their standard errors, are estimated with an upstream model suited to the data (for example, linear or negative-binomial regression, or pseudobulk DESeq2). For each target-outcome pair, PETRO weights guides by their agreement with the guide consensus, shown for target 1 and outcome *Y*_2_: weights decrease with discordance and reach zero for *g*_4_ (bottom), and the blue line marks the PETRO estimate. PETRO returns effect estimates and *P* values for all target-outcome pairs, together with guide weights for each outcome; crosses mark zero weights. **b**, Guides act on outcomes through the intended target (solid arrows) and, when off-target, through paths that bypass it (red dashed arrows). Bottom, *g*_3_ acts on its intended target through a full match and on outcome *Y*_1_ through a partial seed match at the *Y*_1_ promoter. **c**, Outcome-specific guide validity. Top, estimated guide effects of five guides on three outcomes; dashed lines mark the on-target perturbation effect Γ. Bottom, PETRO weights. *g*_3_ deviates only for *Y*_1_ and *g*_4_ only for *Y*_2_, and each receives zero weight only for that outcome. **d**, Population guide effects with zero to three of five guides invalid (red points). Pooling (red diamonds) and PETRO (blue diamonds) both recover the on-target perturbation effect Γ (dashed lines) when all guides are valid, but only PETRO does so when one or two guides are invalid. With three invalid guides, the majority rule is violated and recovery is not guaranteed. **e**, Proportional extension. Left, guides with different effects on target gene expression (*g*_1_ weak, *g*_5_ strong) act on outcome *Y* through the same target. Middle, under the proportional model, valid guides lie on a line through the origin with slope *α*. Right, two-stage least squares (2SLS), which treats all guides as valid instruments, is shifted by invalid guides, whereas PETRO recovers the slope *α* shared by the valid majority.

Under the common on-target perturbation effect model,

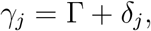

where Γ is the on-target perturbation effect shared by guides targeting the same target and *δ*_*j*_ is the guide-specific off-target perturbation effect (Fig. 1b). Guide *j* is valid for a target-outcome pair when *δ*_*j*_ = 0. Because *δ*_*j*_ is outcome-specific, the same guide can be valid for one outcome and invalid for another (Fig. 1c).

Without restrictions on the off-target effects, Γ is not identified even if the guide effects are known exactly. Pooling and the equal-guide mean assign fixed weights to guides and therefore recover Γ only when all guides are valid or their off-target effects cancel. Under the majority rule, by contrast, more than half of the guide effects equal Γ, making Γ the unique majority-supported value and the population guide median (Fig. 1d). PETRO estimates this effect by robust M-estimation, downweighting guides whose estimated effects depart from the others using Tukey-bisquare weights [16]. Because weights are computed separately for each outcome, a guide can be downweighted for one outcome and retained for another.

Same-target guides can also differ in how strongly they perturb their intended target. When downstream effects scale with target-gene perturbation, the proportional extension uses

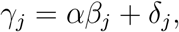

where *β*_*j*_ is the guide’s effect on target gene expression and *α* is the regulatory effect per unit change in target expression (Fig. 1e). A valid guide satisfies *δ*_*j*_ = 0 and therefore follows the common slope even if its target perturbation strength differs from that of other guides. When the analyzed guides have nonzero *β*_*j*_, the majority rule identifies *α*, which PETRO estimates using the same robust weighting of guide-level Wald ratios (Methods).

Two-stage least squares (2SLS) also estimates *α* but treats all guides as valid instruments, so guide-specific off-target effects can shift its estimate away from the slope shared by the valid majority. Canon likewise targets *α* but requires the proportional model and models off-target effects through a spike-and-slab prior [12]; PETRO instead permits arbitrary off-target effects in the invalid minority and applies under either the common-effect or proportional model. This distinction is relevant because held-out-guide prediction showed that proportionality was not uniformly favored across target-outcome pairs (Supplementary Fig. S1, Supplementary Note 2).

### Seed-associated off-target bias persists after pooling cells across guides and is reduced by PETRO

We used PAM-proximal promoter seed matching as an external benchmark of off-target binding risk across CRISPRa and CRISPRi screens [10]. Because seed-match length is determined from guide and promoter sequence, rather than from expression responses, and is not used by PETRO, it provides an independent test of whether PETRO reduces sequence-associated off-target bias (Methods). In the RPE-1 CRISPRa data, the mean guide-level bias increased from 0.024 at 9 bp to 0.194 at 11 bp and 0.460 at 13-18 bp (Fig. 2a). The same qualitative pattern appeared across additional datasets, with longer matches associated with excess activation in CRISPRa and excess repression in CRISPRi (Fig. 2b).

**Fig. 2.**
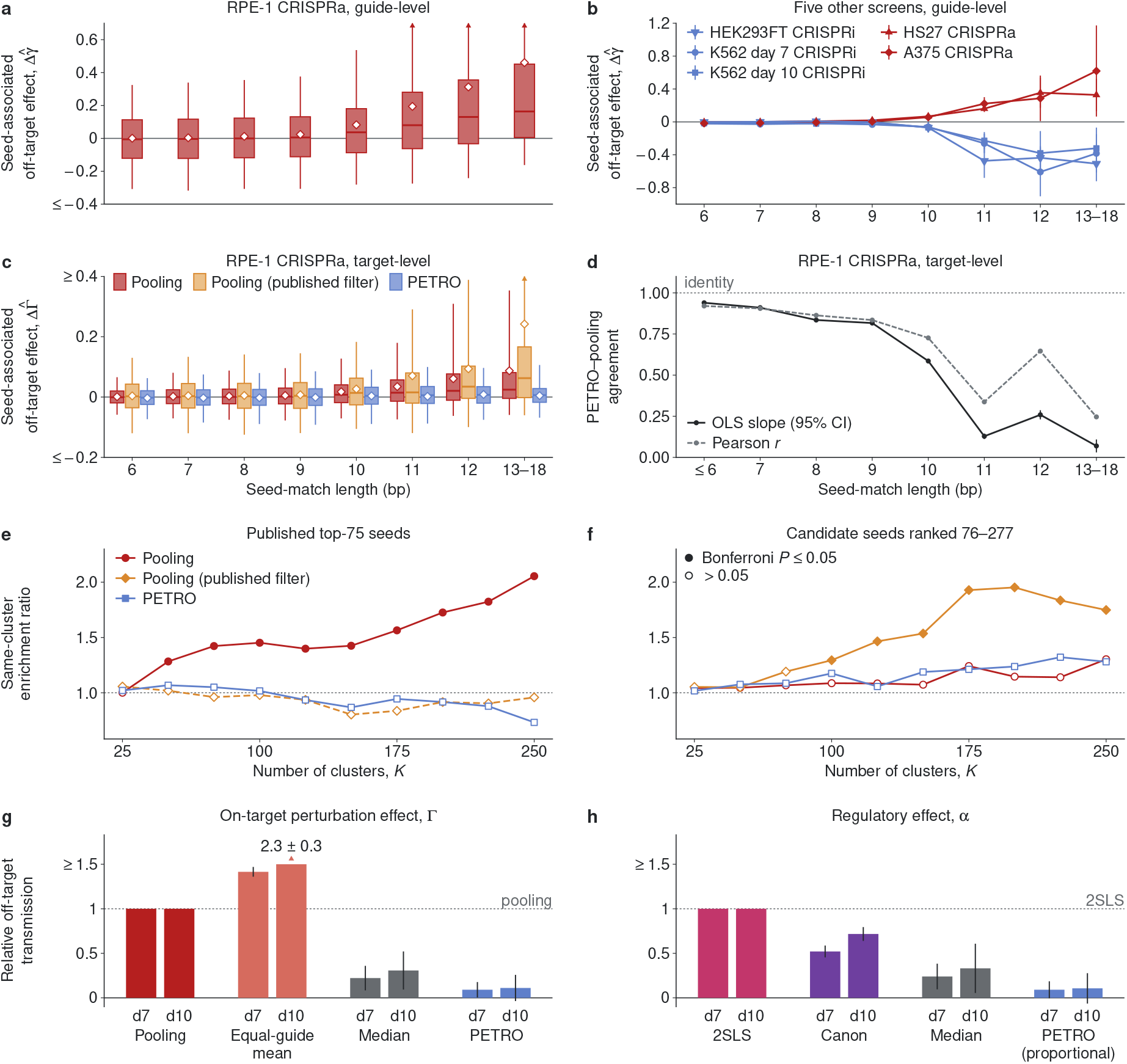
PETRO reduces seed-associated off-target bias and promiscuous-guide transmission in Perturb-seq. **a**, Guide-level seed-associated bias by PAM-proximal guide-promoter seed-match length in RPE-1 CRISPRa. Bias is the estimated effect minus a pooled same-target reference from guides with *<* 6-bp matches to the outcome. **b**, Mean guide-level bias across additional CRISPRa and CRISPRi datasets. **c**, Target-level bias under pooling, pooling after the published filter and PETRO on unfiltered guides. **d**, Ordinary least-squares (OLS) slopes (PETRO on pooling, with an intercept) and Pearson correlations between PETRO and pooling point estimates. In **c**,**d**, target-outcome pairs are grouped by their longest guide-promoter seed match. **e**,**f**, Same-cluster enrichment among RPE-1 targets linked by the 75 seeds selected by the published filter (**e**) and candidate seeds ranked 76-277 (**f**). Seed links were defined from pre-filter (**e**) or retained (**f**) guide sets and held fixed across estimators. Enrichment is relative to the permutation expectation. Filled symbols indicate Bonferroni-adjusted one-sided permutation *P* ≤ 0.05 across ten cluster counts *K*; open symbols indicate adjusted *P >* 0.05. **g**,**h**, Relative transmission of externally flagged promiscuous-guide deviations into estimates of the on-target perturbation effect Γ (**g**) and regulatory effect *α* (**h**) in K562 CRISPRi at days 7 and 10. Transmission is evaluated across outcomes with SCEPTRE *q <* 0.05 for the flagged guide and normalized to pooling (**g**) or 2SLS (**h**) = 1 (Methods). In **a**,**c**, boxes show medians and interquartile ranges, whiskers the 5th-95th percentiles and diamonds means. Error bars show guide-clustered 95% confidence intervals (**b**), 95% confidence intervals for slopes (**d**) and ±1 s.e. from a leave-one-target-out jackknife (**g**,**h**; six targets at day 7, seven at day 10). Triangles mark truncated whiskers (**a**,**c**) or bars capped at 1.5 (**g**,**h**); capped bars are labeled with their actual value ± s.e.

Pooling cells across guides attenuated but did not eliminate this signal. For RPE-1 target– outcome pairs whose longest guide match was 13–18 bp, mean target-level bias was 0.087, approximately 19% of the guide-level bias and close to the longest-matching guide’s mean pooling weight of 18% (Fig. 2c). Thus, pooling largely diluted the guide-specific bias according to the carrier guide’s contribution rather than removing it.

Post hoc guide filtering also did not eliminate the independently defined bias. Among 1,017 target–outcome pairs estimable both before and after the published filter, mean bias across the 11–18-bp strata increased from 0.048 to 0.089 (Fig. 2c). This increase is consistent with renormalization of the surviving guides when other same-target guides are removed.

By contrast, PETRO applied to the unfiltered guide effects kept mean target-level bias near zero, with residual bias of 0.005 at 13-18 bp in RPE-1 (Fig. 2c), with similar reductions in other CRISPRa and CRISPRi datasets (Supplementary Fig. S2). Agreement between PETRO and pooling was high at low seed-match risk but declined sharply with longer matches (Fig. 2d), indicating that PETRO made its largest adjustments where the independent sequence benchmark indicated the greatest off-target risk.

### PETRO suppresses seed-driven off-target structure within and beyond the filtering cutoff

The published Southard et al. filter was designed to remove guides whose transcriptional profiles clustered according to shared PAM-proximal seed sequences rather than intended target [3]. We therefore asked whether filtering and PETRO removed this seed-driven structure among the 314 RPE-1 targets, considering both the top 75 ranked seeds used to flag guides for removal and lower-ranked candidate seeds that remained after filtering (Methods).

For the published top-75 seeds, pooling before filtering showed strong enrichment of same-seed target pairs within transcriptional clusters (*P*_any,*K*_ = 6.25 × 10^−5^), whereas enrichment was not significant after filtering or under PETRO (*P*_any,*K*_ = 0.159 and 0.303, respectively; Fig. 2e). Thus, the published filter suppressed the structure associated with the seeds it selected, and PETRO achieved comparable suppression without using the seed ranking or removing guides globally.

The pattern reversed for candidate seeds ranked 76–277. Same-seed enrichment was significant after filtering (*P*_any,*K*_ = 6.25 × 10^−5^) but not under pooling before filtering or under PETRO (*P*_any,*K*_ = 0.240 and 0.208, respectively; Fig. 2f). This is consistent with the renormalization observed in the promoter-matching analysis: removing some guides can increase the relative contribution of surviving guides carrying other sequence-associated effects.

Extending the filter to these lower-ranked seeds would substantially reduce coverage: removing their carriers would discard 59.8% of the guides retained by the published filter and leave 73 of 314 targets without an eligible guide. Thus, PETRO matched the filter’s suppression of structure associated with the selected seeds while avoiding the significant lower-ranked seed enrichment observed after filtering and retaining the full eligible guide set.

### PETRO reduces transmission of off-target effects from externally flagged promiscuous guides

As a third external benchmark, we used seven promiscuous CRISPRi guides in K562 cells independently flagged by Drepanos et al. based on reproducible guide-specific transcriptional effects across days 7 and 10 and supported by comparison with CRISPR knockout perturbations [9]. We analyzed each time point separately, retained the flagged guides during estimation and used their labels only for evaluation. Six flagged guides were evaluable at day 7 and all seven at day 10 (Methods).

For each flagged guide, we measured how much of its deviation from the eligible unflagged same-target guides was transmitted into the corresponding target-level estimate across outcomes significantly affected by that guide (Methods). For the on-target perturbation effect Γ, transmission under PETRO was 0.091 at day 7 and 0.111 at day 10 relative to pooling, which was normalized to 1 (Fig. 2g). Thus, PETRO reduced transmission of these independently flagged guide-specific effects by about 90% without using the flags during fitting.

We next evaluated the regulatory effect *α*, placing the proportional extension of PETRO on the same estimand as 2SLS and Canon [12]. Canon reduced transmission relative to 2SLS to 0.521 at day 7 and 0.717 at day 10, whereas the proportional extension of PETRO transmitted 0.091 and 0.106, respectively (Fig. 2h). At the individual-guide level, transmission was lower under PETRO than under Canon for all evaluable flagged guides: six of six at day 7 and seven of seven at day 10 (two-sided exact sign tests, *P* = 0.0313 and *P* = 0.0156, respectively).

Thus, independently flagged off-target effects propagated into conventional estimates of both Γ and *α*. Majority-based estimation substantially reduced their contribution while retaining the flagged guides and making no use of the external annotations.

### PETRO reduces off-target bias and controls false rejection of the target null

We simulated four-guide targets with three valid guides and one guide carrying an off-target perturbation effect, so the majority rule held but the invalid guide was unknown to the estimators (Methods).

With a strong off-target effect (*δ* = 0.80) and 150 cells per guide, pooling had bias 0.20, whereas at Γ = 0 the mean bias was 0.035 for the guide median and 5.37 × 10^−4^ for PETRO. Across Γ = 0, 0.10 and 0.20, PETRO closely matched the oracle estimator that used only the three valid guides (Fig. 3a).

**Fig. 3.**
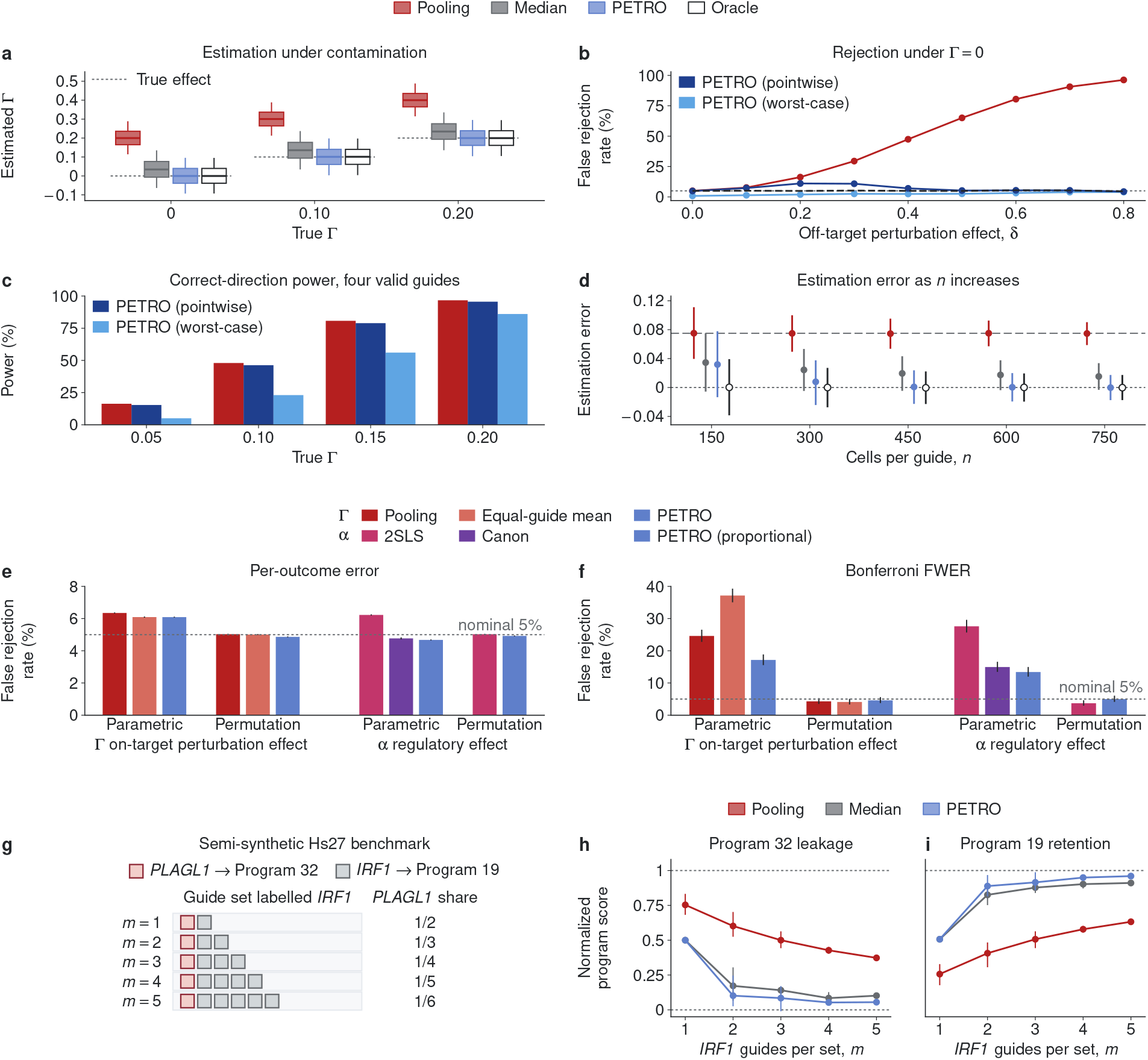
Evaluation of PETRO estimation, inference and biological-signal preservation. **a-d**, Four-guide simulations with 10,000 replicates per condition. The first three guides have guide effect Γ and the fourth has guide effect Γ + *δ*, with residual variance 1. Pooling, the guide median and PETRO use all four guides; the oracle uses the first three. **a**, Estimated on-target perturbation effect Γ for Γ = 0, 0.10 and 0.20, with *δ* = 0.80 and *n* = 150 cells per guide. Boxes show medians and interquartile ranges; whiskers span the 5th-95th percentiles. **b**, False rejection under Γ = 0 as the off-target perturbation effect *δ* increases, with 150 cells per guide. **c**, Correct-direction power, defined as rejection with the correct estimated sign, when all four guides are valid and each has 150 cells. In **b**,**c**, tests are two-sided at the 5% level, and PETRO pointwise and worst-case tests are shown separately. **d**, Mean bias as cells per guide increase from 150 to 750, with Γ = 0 and *δ* = 0.30. **e**,**f**, Per-outcome false rejection rates (**e**) and Bonferroni familywise error rates across outcomes within each target (**f**), evaluated over 2,000 empirical-null target families constructed from Southard RPE-1 CRISPRa control profiles. Downstream guide effects are null, while effects on target gene expression are preserved. For the regulatory effect *α*, the proportional extension of PETRO is shown. Parametric and permutation inference are compared where available; Canon is shown with its native inference. Error bars show 95% confidence intervals. Dotted lines in **b**,**e**,**f** indicate the nominal 5% level. **g**, Semi-synthetic Hs27 benchmark adding one PLAGL1 guide to sets of *m* = 1-5 IRF1 guides and treating each combined set as an IRF1 perturbation. **h**,**i**, Program 32 leakage (**h**) and Program 19 retention (**i**). Leakage is scaled from the corresponding IRF1-only reference (0) to the PLAGL1-only reference (1); retention is the fraction of the corresponding IRF1-only signal preserved. Points show means and bars the minimum-maximum range across all IRF1-guide subsets (5, 10, 10, 5 and 1 subsets for *m* = 1, …, 5).

Under the null Γ = 0, pooling increasingly mistook the off-target effect for a target effect, with false rejection reaching 96.3% at *δ* = 0.80 (Fig. 3b). The PETRO pointwise test substantially reduced this inflation but peaked at 11.1% false rejection at *δ* = 0.20, where contamination was large enough to perturb the fit but difficult to distinguish from sampling variation. The worst-case test remained below 5% throughout (0.8–4.0%). This protection came at a power cost: at Γ = 0.10, correct-direction power was 48.0% for pooling, 46.2% for the pointwise test and 23.1% for the worst-case test; by Γ = 0.20, these increased to 96.6%, 95.6% and 86.0%, respectively (Fig. 3c). We extended these comparisons to five-guide targets with one or two off-target guides, off-target effects of either sign, and Gaussian or negative-binomial outcomes (Supplementary Fig. S3).

Finally, increasing the number of cells sharpened the distinction between persistent aggregation bias and sampling uncertainty. With Γ = 0 and *δ* = 0.30, pooling retained bias near 0.075 as cells per guide increased from 150 to 750, whereas PETRO bias decreased from 0.032 to approximately zero (−1.78 × 10^−4^; Fig. 3d). Thus, greater precision allowed PETRO to recover the majority-supported effect, while pooling converged more precisely to a biased aggregate.

### Permutation inference controls familywise error under empirical nulls

Robustness to off-target perturbation effects does not ensure calibrated inference. We therefore constructed 2,000 empirical-null target families from RPE-1, preserving the observed expression distribution and guide effects on target gene expression while setting all downstream guide effects to zero (Methods).

At the nominal 5% level, marginal false rejection rates were only modestly miscalibrated. For the on-target perturbation effect Γ, parametric rates were 6.3% for pooling, 6.1% for the equal-guide mean and 6.1% for PETRO, whereas permutation inference gave 4.9–5.0%. For the regulatory effect *α*, parametric rates were 6.2% for 2SLS, 4.8% for Canon and 4.7% for the proportional extension of PETRO; permutation rates were 5.0% for 2SLS and 4.9% for PETRO (Fig. 3e).

The differences were much larger in the extreme null tail relevant to familywise testing. With Bonferroni correction across outcomes, parametric familywise error rates (FWERs) for Γ were 24.6% for pooling, 37.2% for the equal-guide mean and 17.2% for PETRO, compared with a nominal 5%. Permutation inference reduced these to 4.1-4.6% (Fig. 3f). For *α*, parametric FWERs were 27.6% for 2SLS and 13.4% for the proportional extension of PETRO, while Canon’s native inference had an FWER of 15.0%. Permutation inference gave FWERs of 3.7% for 2SLS and 5.0% for the proportional extension of PETRO.

Thus, modest marginal miscalibration can produce substantial familywise error inflation. Permutation inference kept both marginal and familywise error close to nominal for every estimator for which it was evaluated, showing that robustness to guide-specific contamination and calibration of inference address distinct sources of error.

### PETRO limits program misattribution while preserving biological signal

Perturb-seq effects are often interpreted through coordinated transcriptional programs, so guide-specific effects can propagate to higher-level biological conclusions. In the Hs27 CRISPRa data, IRF1 was strongly associated with antigen-presentation Program 19 and PLAGL1 with inflammatory Program 32 [3]. We constructed a semi-synthetic benchmark by adding one observed PLAGL1 guide with a strong Program 32 effect to sets of one to five IRF1 guides and treating the combined set as an IRF1 perturbation (Fig. 3g). Program 32 leakage measured misattribution of the PLAGL1 response, whereas Program 19 retention measured preservation of the intended IRF1 response (Methods).

With one IRF1 and one PLAGL1 guide, no strict majority existed and the IRF1 effect was not identified. In this two-guide boundary case, PETRO and the guide median returned the midpoint of the guide effects, giving Program 32 leakage of 0.500, compared with 0.753 under pooling (Fig. 3h). Once two or more IRF1 guides were combined with the PLAGL1 guide, the IRF1 guides formed a strict majority and PETRO progressively suppressed Program 32 leakage while preserving Program 19 (Fig. 3h,i).

With five IRF1 guides, Program 32 leakage was 0.055 under PETRO versus 0.373 under pooling, while Program 19 retention was 0.960 versus 0.632. The guide median gave leakage of 0.102 and retention of 0.911. A corresponding KLF5 benchmark showed the same qualitative pattern (Supplementary Fig. S4a,b).

We finally asked whether this correction preserved reported biological associations in the original Hs27 analysis. Pooling and PETRO agreed for all 13 eligible published target–program associations (Supplementary Fig. S4c). Thus, PETRO reduced program misattribution in the semi-synthetic benchmarks while preserving strong biological associations in the original data.

### PETRO selectively corrects guide-specific bias without global guide removal

A single global label of guide validity cannot capture outcome-specific disagreement. In RPE-1, all five active PRDM5 guides passed the published filter, yet FTL, the largest pooled PRDM5 effect among 5,424 outcomes, had an estimated effect of 0.961 under pooling and −0.079 under PETRO, driven almost entirely by one guide carrying a terminal C|GGCG motif (Fig. 4a). Across 48 targets with terminal N|GGCG carrier guides, carriers showed excess FTL activation relative to same-target noncarriers in 45 of 48 targets in RPE-1 and 40 of 48 in Hs27; 21 of the 48 carriers were retained by the published filter and showed similar excess activation (Fig. 4b). Thus, globally retained guides can still carry recurrent outcome-specific effects.

**Fig. 4.**
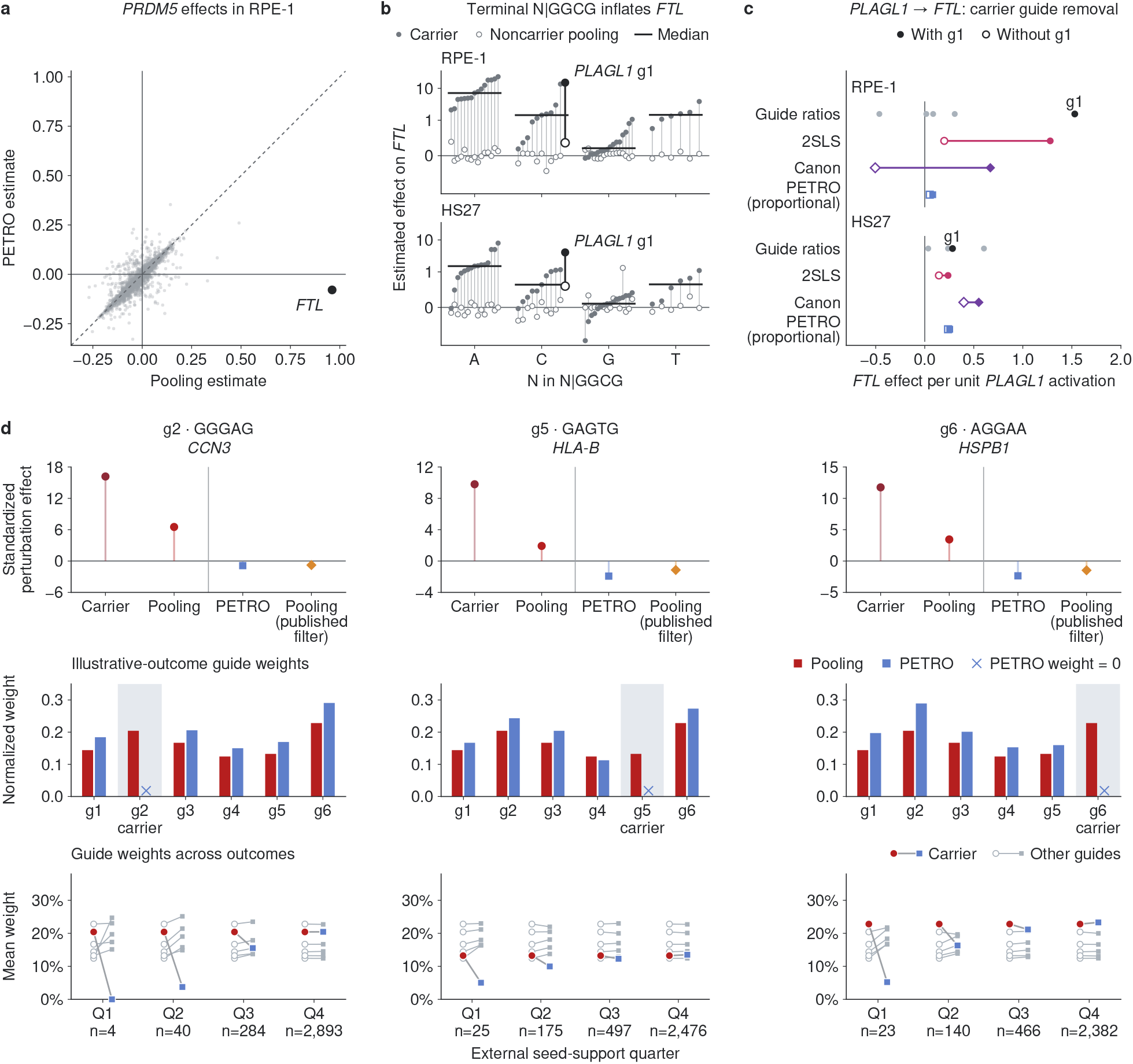
PETRO selectively corrects guide-specific bias without global guide removal. **a**, Pooling and PETRO estimates of the on-target perturbation effect of PRDM5 across 5,424 outcomes in RPE-1 CRISPRa. FTL, the outcome with the largest pooled estimate, is highlighted; the dashed line denotes identity. **b**, Estimated FTL effects of 48 terminal N|GGCG carrier guides in each of RPE-1 and Hs27, grouped by the nucleotide preceding GGCG. Filled points show carrier-guide effects and open points pooled effects from eligible same-target noncarrier guides; vertical lines connect each carrier to its reference. Horizontal bars show carrier-guide medians. PLAGL1 g1 is highlighted. **c**, PLAGL1-FTL regulatory-effect estimates in RPE-1 and Hs27. Guide rows show Wald ratios: each guide’s estimated FTL effect divided by its estimated effect on PLAGL1 expression. Estimator rows compare fits with g1, the C|GGCG carrier (filled symbols), and without g1 (open symbols); lines connect the two estimates. Guide ratios, 2SLS and the proportional extension of PETRO use guides with *P <* 0.05 for their effects on PLAGL1 expression; Canon uses all six guides before removal. Both PETRO fits after removal use the median fallback (Methods). **d**, Outcome-specific weighting of MLXIP guides g2 (GGGAG), g5 (GAGTG) and g6 (AGGAA) in RPE-1, all removed by the published filter. External same-seed profiles were constructed from guides targeting other genes. CCN3 (g2), HLA-B (g5) and HSPB1 (g6) were selected retrospectively as the largest positive contributors to each carrier’s correlation with its external profile (Methods). Top, estimated effects divided by their standard errors for the carrier guide, pooling, PETRO and pooling after filtering. Middle, normalized guide weights; shading marks focal carriers and crosses indicate zero PETRO weight. Bottom, mean guide weights across four outcome groups containing approximately equal shares of total positive correlation contribution, ordered from strongest to weakest support (Q1-Q4). Circles and squares denote pooling and PETRO, respectively; lines connect weights for the same guide, with other guides shown in gray. The displayed *n* values give the numbers of outcomes in each group.

PLAGL1 g1 passed the published filter despite carrying C|GGCG and was the source study’s strongest PLAGL1 activator in both datasets [3]. To test whether its stronger activation explained the FTL excess, we divided FTL effects by PLAGL1 activation; g1 remained strongly discordant in RPE-1 but not in Hs27 (Fig. 4c). Removing g1 changed the RPE-1 2SLS estimate from 1.279 to 0.199 and the Canon estimate from 0.669 to −0.505, whereas the proportional extension of PETRO changed modestly from 0.081 to 0.051. Thus, the RPE-1 carrier exerted substantial leverage on the 2SLS and Canon estimates even after accounting for target activation, whereas the PETRO estimate changed little.

The converse occurred for MLXIP, for which the published filter removed three of six guides. We constructed external transcriptional profiles for their terminal seeds from guides targeting other genes and used these profiles to identify outcomes with strong seed-associated support (Methods). For the strongest illustrative outcomes CCN3, HLA-B and HSPB1, the corresponding carrier guide drove the pooled estimate, whereas PETRO assigned that carrier zero weight and produced an estimate close to pooling after global guide removal (Fig. 4d). Across outcomes, PETRO reduced carrier weight most strongly where external seed support was greatest and allowed it to approach its pooling weight as support weakened. Relative to pooling, PETRO reduced the squared correlation of the MLXIP profile with the external GGGAG, GAGTG and AGGAA profiles by 80.3%, 43.2% and 65.6%, respectively. Thus, guides removed globally can still contribute where they agree with same-target guides while being downweighted selectively where guide-specific effects appear.

MLXIP–S100A6 exposed the limit of guide agreement alone. The three globally flagged guides had the largest effects (0.618–0.966), whereas the other three ranged from −0.416 to 0.294 (Supplementary Fig. S5a). The PETRO estimate (0.344) fell between the groups and rested on only 2.65 effective guides out of six (Methods), so PETRO flagged it for low precision-weighted agreement (Supplementary Fig. S5b,c). Here guide agreement alone does not provide a majority-supported effect, and external sequence evidence can therefore be informative.

### Guide disagreement changes the biological interpretation of ZNF219 perturbation

We finally asked whether guide disagreement could alter the biological interpretation of a perturbation rather than only individual target-outcome estimates. In a human cortical Perturb-seq study, pooled ZNF219 knockdown was reported to reduce inhibitory-lineage representation, deplete inhibitory progenitors, enrich postmitotic inhibitory neurons and increase excitatory relative to inhibitory output, leading to the interpretation that ZNF219 represses neuronal differentiation [11]. All three ZNF219 guides passed the study’s knockdown filter and achieved comparable target knockdown, yet they supported different components of this phenotype (Fig. 5a). Overall inhibitory-lineage depletion was significant only for g3, whereas g1 significantly increased inhibitory-lineage representation. Both g1 and g3 depleted inhibitory progenitors, but only g1 significantly enriched postmitotic inhibitory neurons. Thus, no individual guide reproduced the full pooled pattern: pooling combined inhibitory-lineage depletion supported by g3 with postmitotic enrichment supported by g1. By contrast, guides targeting ARX and NR2E1 were directionally concordant with their pooled estimates.

**Fig. 5.**
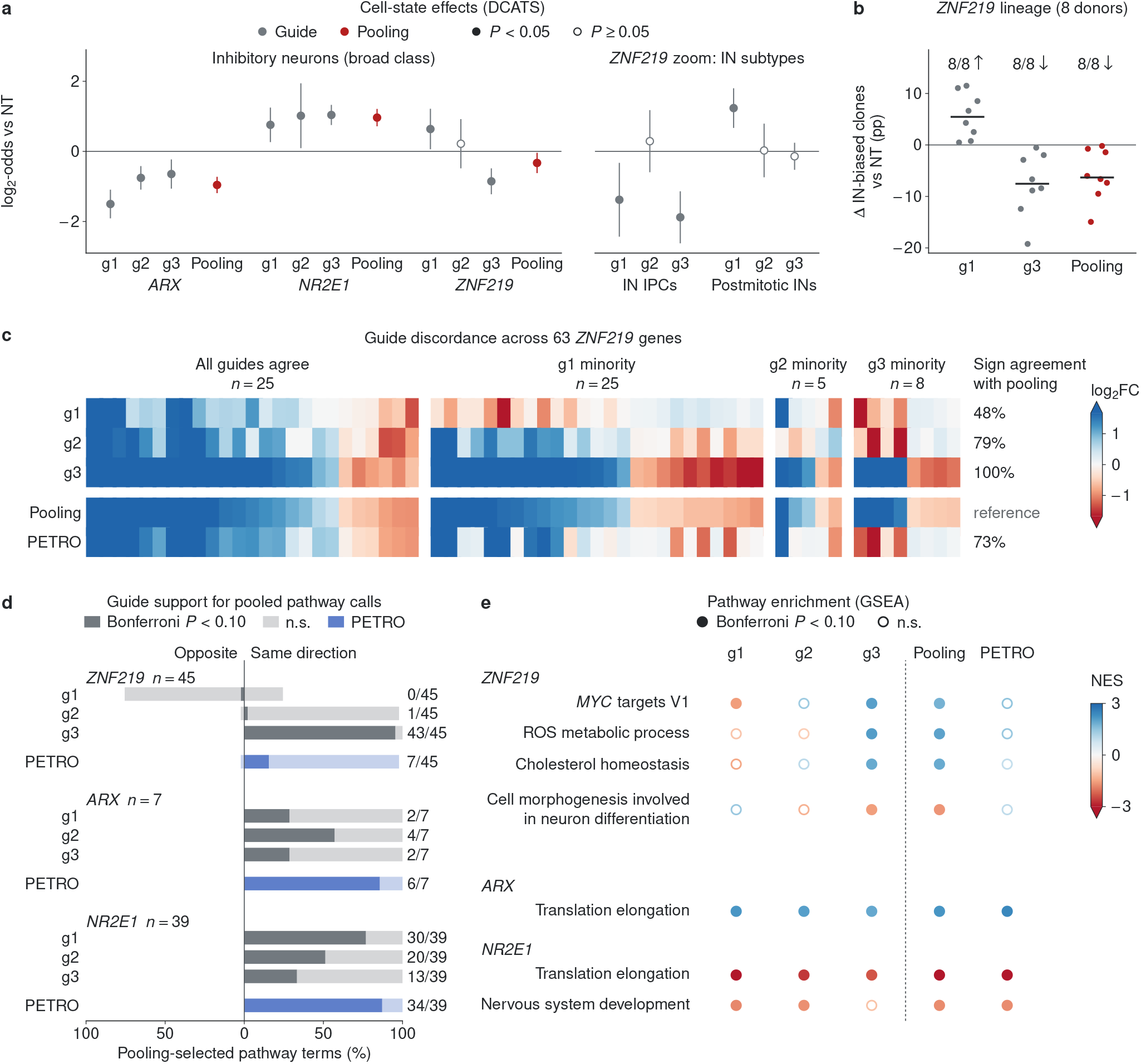
Guide pooling combines conflicting ZNF219 effects. Reanalysis of cortical Perturb-seq data from Ding et al. [11]. **a**, DCATS log_2_ odds ratios relative to non-targeting controls (NT), estimated from day-7 cells from four donors, for the overall inhibitory lineage and, for ZNF219, inhibitory progenitors (IN IPCs) and postmitotic inhibitory neurons. Gray and red circles show guide-level and pooled estimates, respectively. Open symbols indicate unadjusted DCATS likelihood-ratio *P* ≥ 0.05; error bars show model-based 95% Wald confidence intervals. **b**, Changes in inhibitory-neuron-biased clone fractions relative to NT, in percentage points, across eight donors. Points represent individual donors; bars show medians and annotations give the number of donors with the indicated direction. **c**, Guide-level, pooled and PETRO log_2_ fold changes for the 63 genes in the published ZNF219 inhibitory-neuron network, grouped by whether all three guides agree in sign or g1, g2 or g3 opposes the other two. Percentages show agreement in sign with pooling across all 63 genes. **d**, Guide and PETRO support for pathway terms significant under pooling. Bars show the percentages of these terms enriched opposite to (left) or in the same direction as (right) pooling. Darker segments indicate significant enrichment. Fractions give the number of significant same-direction terms divided by the number of pooling-significant terms for each target; individual terms may be supported by multiple guides. **e**, Selected gene set enrichment analysis (GSEA) results for ZNF219 and the concordant ARX and NR2E1 controls. Colors show normalized enrichment scores; filled and open symbols denote significant and nonsignificant enrichment, respectively. In **d**,**e**, significance is Bonferroni-adjusted *P <* 0.10, with correction within each target, analysis and gene-set collection before pathway selection.

The opposing ZNF219 effects recurred in a lineage-tracing experiment from the same study. In all eight donors, g1 increased the fraction of inhibitory-neuron-biased clones, whereas g3 and pooling decreased it (Fig. 5b). This consistency across donors strengthened the evidence for opposing guide-specific phenotypes rather than a common ZNF219 effect.

Guide disagreement also extended to the transcriptional response. Among the 63 genes in the published ZNF219 inhibitory-neuron network, the pooled profile matched g3 in sign at all 63 genes, compared with 47.6% agreement for g1 and 79.4% for g2 (Fig. 5c). All three guides agreed at 25 genes; among the remaining 38, g1 was the minority guide at 25, g2 at five and g3 at eight. Thus, the identity of the discordant guide varied across outcomes, directly motivating outcome-specific rather than global guide weighting.

This disagreement propagated to pathway interpretation. Among 45 pathway terms significant for the pooled ZNF219 profile, 43 were also significant in the same direction under g3, compared with one under g2 and none under g1 (Fig. 5d). For several terms, g1 instead showed enrichment in the opposite direction; for example, MYC targets V1 was negatively enriched under g1 and positively enriched under g3. Thus, the pooled pathway profile was supported predominantly by g3 even though other components of the pooled cell-state phenotype were supported by different guides.

We then applied PETRO separately to each gene, requiring majority support while allowing the minority guide to vary across outcomes (Methods). The resulting profile matched the pooled profile in sign at 73.0% of the 63 network genes, but only 7 of the 45 pooling-significant pathway terms retained significant enrichment in the same direction (Fig. 5c,d). This loss was not a general consequence of robust aggregation: 6 of 7 pooling-significant ARX terms and 34 of 39 NR2E1 terms retained same-direction significance under PETRO.

The cause of the ZNF219 disagreement remains unresolved. Comparable knockdown argues against a simple failure to perturb the intended target but does not establish whether the discordance arose from off-target activity or another guide-specific mechanism. Nor do these data identify which individual guide phenotype represents the biological effect of ZNF219. They show instead that the pooled biological interpretation depends strongly on how conflicting replicate guides are combined: requiring majority support substantially changed the ZNF219 profile while preserving most pathway calls for the concordant ARX and NR2E1 controls.

## Discussion

Perturb-seq uses multiple guides as replicates of an intended perturbation, but replication alone does not determine the target-level effect when individual guides carry outcome-specific deviations. Conventional aggregation estimates the on-target perturbation effect plus a weighted combination of these deviations, so increasing the number of cells improves precision without resolving the attribution problem. Under the majority rule, replicate-guide agreement provides identification: if more than half of the guides are valid for a target-outcome pair, the on-target perturbation effect is uniquely identified without knowing which guides form that majority [13, 14]. When no strict majority exists, guide agreement alone cannot identify the effect, as illustrated by the two-guide boundary case of the IRF1 benchmark and by MLXIP–S100A6, which PETRO flagged for low agreement.

Defining validity for each target-outcome pair also clarifies the role of guide filtering. Sequence- and profile-based diagnostics provide valuable evidence about off-target liability [3, 8–10], but global guide removal and target-outcome-specific robust estimation serve different purposes. We found that retained guides can carry large deviations for particular outcomes, whereas removed guides can remain concordant with same-target guides for others. Filtering can also increase the relative influence of surviving guide-specific effects through weight renormalization. Guide annotations and robust target-level estimation are therefore complementary rather than interchangeable.

Robust estimation and calibrated inference are likewise distinct. Uncertainty about the valid-guide set can inflate the false rejection rate of the pointwise PETRO test in finite samples, whereas the worst-case test provides stronger protection at a cost in power. Separately, parametric approximations can be miscalibrated in Perturb-seq data [5]; in our empirical-null analyses, modest marginal miscalibration translated into substantial familywise error inflation. Permutation inference brought both marginal and familywise error rates close to nominal. Thus, robustness to off-target effects does not remove the need for careful calibration of inference.

The primary estimand in PETRO is the on-target perturbation effect Γ, distinct from the target-expression-mediated regulatory effect *α* estimated by instrumental-variable approaches [12, 17, 18]. Recovering *α* additionally requires the proportional model to be appropriate, and held-out-guide prediction showed that proportionality was not uniformly favored across target-outcome pairs. We used the proportional extension of PETRO for direct comparison with Canon. In the promiscuous-guide benchmark, both methods reduced transmission of flagged off-target effects relative to conventional 2SLS, with lower transmission under PETRO for every evaluated guide. These comparisons were conducted in the few-guide regime considered here and do not isolate which modeling differences account for the observed performance.

The consequences of guide disagreement can extend to biological interpretation. In our reanalysis of cortical Perturb-seq data [11], three ZNF219 guides with comparable knockdown supported conflicting cell-state and transcriptional responses. The identity of the discordant guide varied across genes, and the pooled transcriptional profile was dominated by one guide even though other components of the pooled phenotype were supported by another guide. Requiring majority support separately for each outcome substantially changed the ZNF219 profile, while most pathway calls for the concordant ARX and NR2E1 controls were retained. Pooling and PETRO also agreed for all 13 eligible published Hs27 target–program associations. These analyses do not identify which ZNF219 guide phenotype is biologically correct or establish the cause of the disagreement; they show that pooled interpretation can depend strongly on how conflicting replicate guides are combined.

Several limitations define the scope of the framework. The majority rule is an identification assumption and cannot generally be verified from the observed guide effects alone. If at least half of the guides are invalid, PETRO can fail to recover the on-target perturbation effect, particularly when invalid guides agree with one another. In finite samples, small off-target effects may also be difficult to distinguish from sampling noise. In addition, the assumed form of the on-target effect may not fit every target-outcome pair. The common-effect model assumes a shared ontarget perturbation effect across guides, whereas the proportional extension assumes downstream effects scale with target-gene perturbation. More general forms of on-target heterogeneity would require additional structure or information. For genomic-region targets that alter multiple genes, Γ represents the effect of perturbing the region and need not isolate the regulatory effect of any one gene.

Because PETRO operates on guide-level estimates and their uncertainty, it can be applied after linear, count-based or other upstream models. The majority rule also has implications for experimental design: tolerating up to *k* invalid guides requires at least 2*k* + 1 guides per target, so odd guide counts use replication more efficiently under this criterion. More broadly, the majority rule is an identification principle rather than a feature specific to PETRO. When guide-specific effects are possible, replicate guides determine not only how precisely a target-level perturbation effect can be estimated, but whether it can be identified at all.

## Methods

### PETRO model and identification

PETRO is a causal inference framework for estimating on-target perturbation effects from multiple guides designed to perturb an intended target *A*, which can be either a gene or a genomic element. For a target-outcome pair (*A, Y*), let *j* = 1, …, *m* index the guides designed to perturb *A*. For each guide *j, γ*_*j*_ denotes the population causal effect of guide assignment on outcome *Y* relative to non-targeting controls, and *γ*_*j*_ denotes its estimate from an upstream model suited to the data, such as linear or negative-binomial regression. We refer to *γ*_*j*_ as the guide effect. We decompose the guide effect as 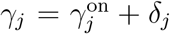, where 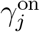 is the on-target perturbation effect operating through perturbation of *A*, and *δ*_*j*_ is the off-target perturbation effect operating through any causal paths that bypass *A*. Off-target paths can arise, for example, when a guide acts at an unintended genomic site through partial seed matches at another promoter [3, 8, 10]. Thus, guides designed to perturb the same target can have different causal effects on a given outcome.

We first consider a common on-target effect model, which assumes that the on-target perturbation effects are the same across guides, 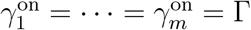. Under this model, Γ is the shared on-target perturbation effect on *Y*, and

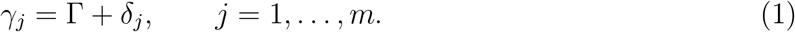

We define a guide as valid for (*A, Y*) when *δ*_*j*_ = 0, and let *V* = {*j* : *δ*_*j*_ = 0} denote the set of valid guides. Because validity is defined separately for each target-outcome pair, the same guide can be valid for one outcome and invalid for another.

Without restrictions on the off-target perturbation effects, Γ is not identified, even if every guide effect *γ*_*j*_ were known exactly, because any candidate value of Γ can be paired with *δ*_*j*_ = *γ*_*j*_ − Γ. We therefore assume the majority rule, under which more than half of the guides are valid (|*V*| *> m/*2). Under this assumption, more than half of the guide effects equal Γ, so Γ is the unique value shared by a strict majority of guides and equals the median of *γ*_1_, …, *γ*_*m*_. The identities of the valid guides do not need to be known, as in the majority rule for invalid instrumental variables [13, 14].

### PETRO estimation

For each target-outcome gene pair, PETRO estimates the on-target perturbation effect Γ while allowing an unknown minority of guides to have arbitrary off-target perturbation effects *δ*_*j*_. Its inputs are the estimated guide effects 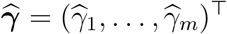 and their estimated covariance matrix 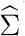 from the upstream model.

For *m* ≥ 3 guides, PETRO chooses baseline weights 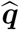 that minimize the estimated variance of the weighted guide average, with each weight capped at *q*_max_ (default 0.4) to limit the contribution of any single guide:

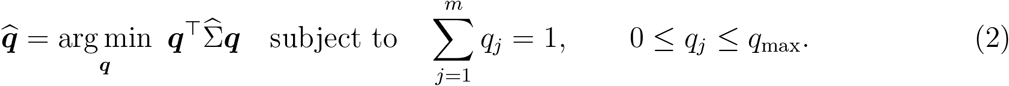

When the estimated guide effects are uncorrelated, these are capped inverse-variance weights. PETRO then downweights guides that disagree with the candidate on-target perturbation effect. For a candidate value *θ* of Γ, define the standardized residual 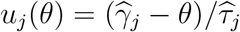, where 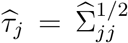 Let 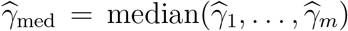. Among the roots of the estimating equation in equation (3), which is based on the Tukey bisquare loss function, PETRO selects the root that is closest to the median:

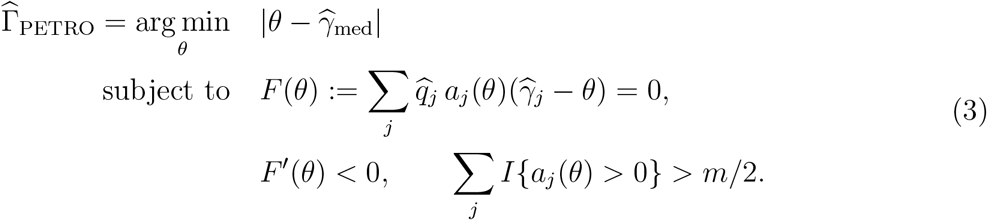

Here, *a*_*j*_(*θ*) = {1−*u*_*j*_(*θ*)^2^*/c*^2^}^2^*I*{|*u*_*j*_(*θ*)| *< c*} is the Tukey bisquare weight with a cutoff *c >* 0 [16]. These constraints make *θ* a Tukey-robust weighted average at a strict local minimum of the loss, supported by more than half of the guides. The Tukey-robust weights decrease from 1 at 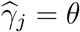 to 0 at |*u*_*j*_(*θ*)| = *c*. PETRO accepts either the conventional fixed cutoff *c* = 4.685 or an adaptive cutoff *c*_n_ with *c*_n_ → ∞ and 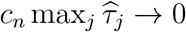. The final weights are 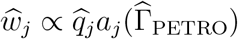, normalized to sum to one. Because 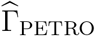 solves the estimating equation, it equals 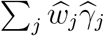. The effective number of guides, 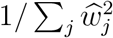, is the number of guides that effectively contribute to the estimate after weighting by precision and agreement; it equals *m* when all guides contribute equally. Estimates with fewer than *m/*2 effective guides are flagged for low precision-weighted agreement. If no *θ* satisfies all constraints, PETRO returns *γ*_med_ and flags the estimate as a fallback. Details of the adaptive cutoff, root search, and fallback are provided in Supplementary Note 1.

### PETRO testing

For each target-outcome pair, PETRO tests the composite null *H*_0_ : Γ = 0 under the majority rule (|*V*| *> m/*2). Let *h* = ⌊*m/*2⌋ + 1 be the smallest strict majority size. For any guide set *S* with |*S*| ≥ *h*, define *H*_0,*S*_ as the null hypothesis that *γ*_*j*_ = 0 for all *j* ∈ *S*, while the effects of guides outside *S* are unrestricted. Because *H*_0_ requires at least *h* valid guides, the composite null can be written as

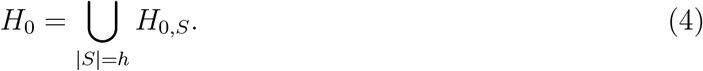

PETRO provides two tests for this null hypothesis. The pointwise test forms a weighted mixture of null distributions for each candidate set *S* where |*S*| ≥ *h*, with weights based on their agreement with 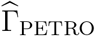. The worst-case test tests all subsets of size *h* and does not require identification of the valid set.

For the pointwise test, the observed test statistic is 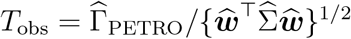, where 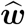 denotes the vector of final PETRO weights and 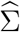 is the estimated covariance matrix of 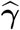.

To generate the null distribution, we first compute mixture weights for all candidate sets *S* with |*S*| ≥ *h*.

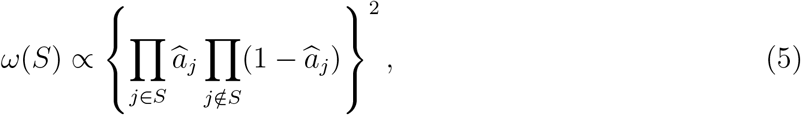

where 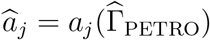. The mixture weights are normalized to sum to one over all candidate sets.

Next, we generate the null distribution of *H*_0,*S*_ by resampling. Specifically for each null replicate *b* = 1, …, *B*, PETRO samples a candidate set *S* according to *ω*(*S*) and generates estimated guide effects under *H*_0,*S*_. A parametric null distribution of *H*_0,*S*_ uses the conditional Gaussian distribution with covariance 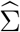. A nonparametric, permutation-based null distribution of *H*_0,*S*_ permutes the labels of guides in *S* and non-targeting controls within prespecified strata, such as batch, while keeping labels for guides outside *S* fixed. The upstream model is refitted after each permutation. Then, PETRO is refitted using all guides and the studentized statistic *T* ^(*b*)^ is calculated. Let 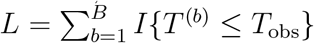 and 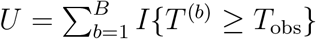 denote the lower- and upper-tail counts among the *B* null replicates. The two-sided Monte Carlo *P* value is *P* = min{1, 2[1 + min(*L, U*)]*/*(*B* + 1)}. Further details are provided in Supplementary Note 1. The worst-case test tests each *H*_0,*S*_ with |*S*| = *h* separately testing the shared effect of the guides in *S*, and obtains a *P* value *p*_*S*_ by parametric or permutation inference. Because *H*_0_ is the union of the subset-specific null hypotheses, the corresponding intersection-union test has *P* value

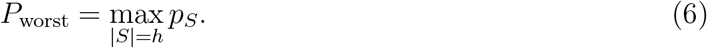

Thus, *H*_0_ is rejected only if every *H*_0,*S*_ is rejected. If each *p*_*S*_ is valid under *H*_0,*S*_, then *P*_worst_ controls type I error. More details are provided in Supplementary Note 1.

### Proportional extension for the regulatory effect

When the intended target *A* is a gene with measured expression, we additionally consider a proportional on-target effect model that allows guides to differ in their on-target perturbation effects according to how strongly they perturb target gene expression. This model replaces the common-effect assumption 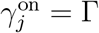 with 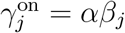, where *β*_*j*_ is the effect of guide *j* on target gene expression and *α* is the regulatory effect, defined as the change in the outcome per unit change in target gene expression on the scale of the guide-level estimates. Combined with the decomposition 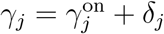, this gives

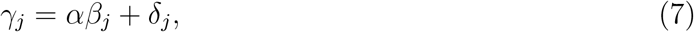

where *δ*_*j*_ remains the off-target perturbation effect. Guide validity therefore remains defined by *δ*_*j*_ = 0. For a valid guide with *β*_*j*_ ≠ 0, *γ*_*j*_*/β*_*j*_ = *α*. Thus, when all analyzed guides have nonzero *β*_*j*_, the majority rule identifies *α* as the ratio shared by a strict majority of guides.

The proportional extension of PETRO estimates *α* from the Wald ratios 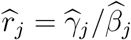, using the same weighting and root-selection procedure as for the common-effect model. In applications, we restrict the proportional analysis to guides with evidence of perturbation of the intended target in the expected direction, as described for each dataset below. This restriction avoids unstable Wald ratios from guides with estimated target-expression effects near zero. The covariance matrix of the Wald ratios is obtained from the joint covariance matrix of the estimated effects on target gene expression and on the outcome using the delta method. Tests of *H*_0_ : *α* = 0 use the same pointwise and worst-case tests. For each candidate valid set *S*, the null restriction is *γ*_*j*_ = 0 for all *j* ∈ *S*, while the guides’ effects on target gene expression remain unrestricted. Further details are provided in Supplementary Note 1.

### Comparator methods

For estimation of the on-target perturbation effect Γ, we compared PETRO with four estimators: (1) pooling, in which all cells carrying guides targeting the same gene were assigned a common perturbation label; (2) pooling after the published filter, which applied pooling only to guides retained by the source study’s guide filter [3]; (3) the equal-guide mean, defined as the unweighted mean of the estimated guide effects; and (4) the guide median, defined as their median.

For estimation of the regulatory effect *α* under the proportional model, we compared the proportional extension of PETRO with three estimators: (1) two-stage least squares (2SLS), using the target’s guide-assignment indicators as instruments (R package ivreg v.0.6-8 [19]); (2) the median of the Wald ratios 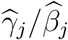 and (3) Canon [12], a one-sample Bayesian instrumental-variable method with spike-and-slab priors on the guides’ effects on target gene expression and on their off-target perturbation effects (R package Canon v.1.0, default settings).

### RPE-1, Hs27, and A375 CRISPRa data

We obtained processed RPE-1 and Hs27 CRISPR activation (CRISPRa) Perturb-seq data from Southard et al. [3] from Zenodo records 15213619 and 15200179. We retained 5,424 RPE-1 and 4,914 Hs27 outcome genes and re-estimated guide effects using the multiple linear regression procedure from the source study.

We retained guides with intended-target activation of at least 0.1 and one-sided Benjamini-Hochberg *q* ≤ 0.05 [20], adjusted across all intended guide-target tests. We analyzed 322 RPE-1 targets with at least five retained guides and 341 Hs27 targets with at least four retained guides. The RPE-1 promoter seed matching, seed enrichment, and PRDM5 and MLXIP case studies were further restricted to the 314 targets with at least one guide retained by the published off-target filter. Guide annotations, published filter calls, the 75 bad seeds and program loadings were obtained from Supplementary Tables 4, 5 and 8 of the source study.

For A375 CRISPRa, we used Akana et al.’s monoculture data [21], retaining guides with positive target-gene effects and *P <* 0.05 in their per-guide test. Targets required at least three retained guides and 20 cells across passing guides. Guide effects were estimated on the log_2_ scale from guide-by-batch pseudobulk counts using edgeR quasi-likelihood models [22] with TMM normalization and batch adjustment.

### HEK293FT and K562 CRISPRi data

For the HEK293FT CRISPR interference (CRISPRi) screen of Kowalski et al. [23], we used the study’s filtered cells and single-guide assignments and retained guides assigned to at least 25 cells. Guide effects were estimated using negative-binomial log-link models without a library-size offset. Models adjusted for replicate, lane and standardized log(1 + *x*) transforms of total UMI count and detected-gene count, each computed without the outcome gene. Each outcome’s dispersion was estimated from control cells and held fixed during guide-effect fitting. Intended-target knockdown was tested using 1,000 one-sided permutations within replicate-by-lane strata. After Benjamini-Hochberg adjustment over the 127 successfully fitted guides, we analyzed the 24 targets with at least four guides having a negative effect on expression of the intended target with *q* ≤ 0.05. Guide effects are reported on the log_2_ scale.

For the K562 CRISPRi data of Drepanos et al. [9], we analyzed days 7 and 10 separately. We analyzed 98 unambiguously assigned CRISPRi targeting guides and ten intergenic control guides from the 25-gene library. Seed-bias analyses used the same negative-binomial model as for HEK293FT, without replicate or lane covariates. For the transmission analysis, we examined the seven guides identified as promiscuous by the source study and used continuous expression to match Canon’s normalized-expression model. Expression was transformed as log(1+counts per 10,000). Expression and guide indicators were then residualized on an intercept, log library size and log detected-gene count, standardized, and used in linear regressions to estimate guide effects on the intended target and outcome.

### Guide seed matching and seed-associated bias

We used PAM-proximal guide-promoter seed matching as an external sequence-based benchmark of off-target binding risk. Following Hartman *et al*. [10], we searched both strands of hg38 within 2 kb of each outcome gene’s annotated transcription start sites for exact matches to the guide’s 3^′^ end adjacent to an NGG protospacer-adjacent motif and recorded the longest match for each guide-outcome pair. Matches shorter than 6 bp were treated as no match, and matches to the intended target and matches of 19 bp or longer were excluded.

The primary RPE-1 analysis used the 314 targets with five or six guides showing evidence of target activation and at least one guide retained by the published filter [3]. For a guide *j* targeting gene *A* and outcome *k*, we defined its seed-associated deviation as

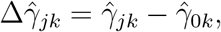

where 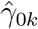 is the pooled effect of same-target guides with a seed match shorter than 6 bp to outcome *k*. Guide-outcome pairs without such a low-match reference were excluded. Guide-level deviations were summarized by seed-match length.

For target-level analyses, pooling, pooling after the published filter and PETRO were compared with the same low-match reference. Target-outcome pairs were grouped according to the longest promoter seed match among the target’s analyzed guides. Comparisons of pooling before and after filtering were restricted to target-outcome pairs estimable under both guide sets. To assess whether PETRO selectively altered estimates associated with greater seed-match risk, we computed, within each seed-match stratum, the Pearson correlation between PETRO and pooling estimates and the ordinary least-squares slope from regressing the PETRO estimates on the pooling estimates with an intercept.

The same analysis was repeated in the additional CRISPRa and CRISPRi datasets using the guide-selection and guide-level outcome models described above.

### Seed enrichment in target clusters

To assess seed-associated structure in target-level transcriptional profiles, we analyzed the same 314 RPE-1 targets under pooling before filtering, pooling after the published filter and PETRO applied to the unfiltered eligible guides. For each target, the same set of target-outcome positions, defined by within-target Benjamini–Hochberg *q <* 0.05 under at least one estimator, was used across estimators.

We considered two sets of PAM-proximal seeds. The first consisted of the 75 cluster-driving seeds selected by the published Southard *et al*. filter [3]. To evaluate whether filtering removed the structure associated with these seeds, same-seed target pairs were defined using the original pre-filter guide sets and held fixed across estimators. Thus, removal of a seed-carrying guide did not itself remove the corresponding target pair from the benchmark.

We additionally reconstructed the source study’s candidate-seed ranking and considered seeds ranked 76-277, which fell below the published top-75 filtering cutoff. For these lower-ranked seeds, same-seed target pairs were defined using guides retained by the published filter and again held fixed across estimators. This construction asks whether seed-associated structure remained among links that survived filtering rather than redefining the benchmark separately for each estimator.

For each estimator, target transcriptional profiles were clustered by average linkage using one minus Pearson correlation as the distance, and the resulting trees were cut at *K* = 25, 50, …, 250 clusters. At each *K*, we counted the number of same-seed target pairs assigned to the same cluster and calculated enrichment relative to its permutation expectation.

To preserve differences in the opportunity for seed sharing across targets, seed annotations were permuted among targets within strata defined by guide count and seed burden while target clusters were held fixed. The same permutations were used across clustering resolutions and estimators. Pointwise permutation *P* values were adjusted across the ten values of *K*. We additionally used a synchronized any-*K* test, *P*_any,*K*_, based on the maximum standardized enrichment across clustering resolutions, with its null distribution obtained from the corresponding maximum in each permutation.

To assess the coverage cost of extending the published filter, we also removed guides carrying candidate seeds ranked 76-277 and recorded the fraction of otherwise retained guides removed and the number of targets left without an eligible guide.

### Transmission of promiscuous-guide deviations

We used the seven promiscuous K562 CRISPRi guides identified by Drepanos *et al*. [9], one for each of seven target genes. The source study defined guide promiscuity using transcriptional profiles from days 7 and 10, which arose from separate transductions. For each guide, its median correlation with other same-target guides was subtracted from its self-correlation across the two time points, and guides with a standardized difference greater than 2 were flagged. The source study further reported that most flagged guides perturbed more genes than their same-target counterparts and that removing them improved agreement between CRISPRi and CRISPR knockout perturbations. We used these guide labels only for evaluation and did not provide them to any estimator.

Days 7 and 10 were analyzed separately. Using SCEPTRE [5], we retained guides with a negative effect on target gene expression (one-sided *P <* 0.05), and targets were required to have at least three retained guides, including the promiscuous guide. Six of the seven flagged guides met these criteria at day 7 and all seven at day 10. Outcomes were selected as those with *q <* 0.05 for the promiscuous guide in the SCEPTRE results released by the source study.

For each evaluated target-outcome pair *i*, a reference *b*_*i*_ was computed from the eligible unflagged same-target guides using reference estimator *ℓ*_0_, where *ℓ*_0_ was pooling for the on-target perturbation effect Γ and 2SLS for the regulatory effect *α*. We defined

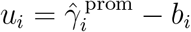

as the promiscuous guide’s deviation from this reference and

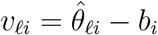

as the corresponding deviation of estimator *ℓ*. For *α*, the promiscuous-guide and estimator deviations were calculated on the Wald-ratio scale and multiplied by the promiscuous guide’s estimated effect on target gene expression so that transmission was expressed on the outcome-effect scale.

Transmission relative to the conventional reference estimator was

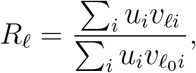

so that 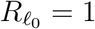, whereas *R*_*ℓ*_ = 0 indicates no net transmission of the promiscuous-guide deviation into estimator *ℓ* across evaluated target-outcome pairs. Standard errors of *R*_*ℓ*_ were estimated using a delete-one-target jackknife and are shown as error bars in Fig. 2g,h.

For the comparison between Canon and the proportional extension of PETRO, we additionally calculated transmission separately for each flagged guide by applying the same definition across that guide’s evaluated outcomes. We compared the paired guide-specific transmission values using a two-sided exact sign test within each time point.

### Simulation study

We simulated a target with four guides, three valid and one with an off-target perturbation effect. Each guide was assigned to *n* cells and all guide groups shared 6*n* non-targeting control cells. Outcomes were generated as *N* (0, 1) in control cells and *N* (Γ + *δ*_*j*_, 1) in cells carrying guide *j*, with *δ*_1_ = *δ*_2_ = *δ*_3_ = 0 and *δ*_4_ = *δ*. Thus, the majority rule held in every contaminated setting, but the identity of the invalid guide was not supplied to the estimators. We generated 10,000 datasets per condition.

We considered four settings: (1) Γ ∈ {0, 0.10, 0.20}, *δ* = 0.80 and *n* = 150 to evaluate point estimation under strong contamination; (2) Γ = 0, *δ* = 0, 0.10, …, 0.80 and *n* = 150 to evaluate false rejection; (3) Γ ∈ {0.05, 0.10, 0.15, 0.20}, *δ* = 0 and *n* = 150 to evaluate power when all guides were valid; and (4) Γ = 0, *δ* = 0.30 and *n* = 150, 300, …, 750 to evaluate estimation as guide-level precision increased. Because guide groups were equally sized, pooling gives equal weight to their effects in this Gaussian setting and has expected bias *δ/*4 when one guide is contaminated.

We compared pooling, the guide median, PETRO and an oracle estimator that combined only the three valid guides using minimum-variance weights. For testing, pooling used a Gaussian cell-level bootstrap under the null, the oracle used correlated Gaussian reference draws, and PETRO used the pointwise and worst-case tests with parametric inference described above. All tests were two-sided at level 0.05. Bias was the mean estimated effect minus Γ; false rejection was the proportion of replicates with *P* ≤ 0.05 when Γ = 0; and correct-direction power was the proportion with *P* ≤ 0.05 and an estimated effect with the true sign.

### Empirical null calibration

To assess calibration under a null distribution that preserves the expression characteristics of the RPE-1 data, we constructed 2,000 empirical-null target families. Fifty targets were selected as templates, 25 with five retained guides and 25 with six, to span variation in effects on target gene expression, cell counts and numbers of cells per droplet without using downstream expression effects. For each null family, one template target was sampled at random.

Every cell retained its original guide assignment but received the full expression profile of a non-targeting control cell sampled with replacement from controls with the same GEM group and recorded number of cells per droplet. We then restored each template guide’s estimated effect on expression of its intended target while leaving all other genes unchanged. Thus, guide effects on target gene expression were preserved, allowing evaluation of the regulatory effect *α*, while the downstream effects satisfied Γ = 0 and *α* = 0 for every tested outcome.

For the on-target perturbation effect Γ, we evaluated pooling, the equal-guide mean and PETRO. For the regulatory effect *α*, we evaluated 2SLS, Canon and the proportional extension of PETRO. Parametric inference used Wald tests for pooling and the equal-guide mean, the *t*-test for 2SLS and Gaussian reference draws for PETRO and its proportional extension (Supplementary Note 1). Canon was evaluated using its published inference procedure.

For permutation inference, guide labels were permuted among cells carrying the target’s guides and non-targeting controls within strata defined by GEM group and recorded number of cells per droplet, and the corresponding guide-level model and target-level estimator were refitted to each permuted dataset. For PETRO and its proportional extension, permutation was restricted to the guides in each candidate valid set as described above.

The marginal false rejection rate was the proportion of null target-outcome tests with *P* ≤ 0.05. To assess calibration in the more extreme tail relevant to genome-wide outcome testing, we applied Bonferroni correction across all outcomes within each target family and defined the FWER as the proportion of the 2,000 families containing at least one rejection.

### Hs27 program analyses

To assess transmission of a guide-specific effect into transcriptional programs, we constructed a semi-synthetic benchmark from the five Hs27 IRF1 guides with at least 20 cells each. IRF1 was associated with Program 19, annotated for antigen presentation, whereas PLAGL1 was associated with the distinct inflammatory Program 32 in the source study [3]. We added PLAGL1 g1, which had a strong Program 32 effect, to each nonempty subset of the five IRF1 guides and treated the resulting guide set as an IRF1 perturbation. This gave 31 IRF1 subsets containing one to five guides, each analyzed alone (clean) and after addition of PLAGL1 g1 (contaminated).

For each of the 206 genes with nonzero loadings on Program 19 or Program 32, we aggregated guide effects using pooling, the guide median and PETRO. With one or two guides, PETRO returned the guide median. For program *k*, the score was

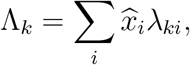

where 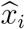 was the aggregated effect on gene *i* and *λ*_*ki*_ was its published program loading.

Program 32 leakage and Program 19 retention were defined as

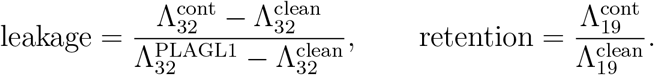

Thus, leakage of 0 corresponds to the IRF1-only reference and leakage of 1 to the PLAGL1-only reference, whereas retention measures the fraction of the corresponding IRF1-only Program 19 signal preserved. For each IRF1 subset size, we report the mean, minimum and maximum across all subsets of that size.

We repeated the benchmark with KLF5 in place of IRF1 and its associated Program 31 in place of Program 19. To assess preservation of previously reported biology in the original Hs27 data, using the published program loadings, we compared pooled and PETRO program scores for 13 eligible target–program associations stated in the main text of the source study, requiring at least three guides with significant activation of the target gene per target.

### Motif carrier and outcome-specific guide analyses

For PRDM5, we compared pooled and PETRO estimates from its five guides across 5,424 RPE-1 outcomes. FTL was selected retrospectively because it had the largest pooled effect and the largest difference between the two estimates. We then asked whether guides ending in GGCG also showed excess effects on FTL across targets. In RPE-1 and Hs27, we identified 48 targets with a terminal N|GGCG carrier guide and compared the carrier’s FTL effect with the pooled effect of its same-target noncarrier guides, excluding noncarriers with published seed annotations [3]. Carriers were grouped by the nucleotide immediately preceding GGCG.

For PLAGL1, we evaluated whether the FTL effect of g1, its terminal C|GGCG carrier, could be explained by its stronger effect on PLAGL1 expression. Guide-specific Wald ratios were calculated as the estimated FTL effect divided by the estimated effect on PLAGL1 expression. We estimated the PLAGL1-FTL regulatory effect before and after removing g1 using 2SLS, Canon and the proportional extension of PETRO. 2SLS and the proportional extension of PETRO used guides whose estimated effect on PLAGL1 expression had *P <* 0.05; Canon used all available guides because it selects guide relevance internally.

For MLXIP, we examined the three guides removed by the published filter, whose protospacers end in GGGAG, GAGTG, and AGGAA. For each 5-mer, we constructed an external profile *e*_*k*_ over outcomes *k* using guides targeting genes other than MLXIP. Among targets with at least one guide ending in the 5-mer and at least two other guides, we calculated the median carrier effect minus the median noncarrier effect for each outcome and then took the median of these differences across targets. For each MLXIP carrier, we computed the Pearson correlation between its effects *x*_*k*_ and the corresponding external profile *e*_*k*_ across the *p* non-MLXIP outcomes. Each outcome’s contribution to this correlation was

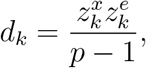

where 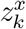 and 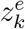 are the standardized carrier and external-profile effects. The outcome with the largest positive *d*_*k*_ for each carrier was selected retrospectively for illustration in Fig. 4d.

Outcomes with *d*_*k*_ *>* 0 were divided into four groups, each accounting for approximately one quarter of the total positive correlation contribution, and we compared the mean weights assigned by pooling and PETRO within each group. Attenuation of seed-associated structure was summarized as

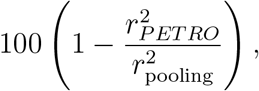

where *r*_*PETRO*_ and *r*_pooling_ are the Pearson correlations of the corresponding MLXIP target-level profiles with the external same-seed profile across all non-MLXIP outcomes.

For MLXIP–S100A6, we assessed whether the six estimated guide effects supported a common effect among a strict majority of guides. Because a strict majority of six requires at least four guides, we considered all 15 four-guide subsets and tested equality of the four guide coefficients using *F* tests from the RPE-1 linear model. Guide weights and the effective number of guides are shown in Supplementary Fig. S5.

### Cortical Perturb-seq analysis

We reanalyzed the cortical CRISPRi Perturb-seq and STICR lineage data of Ding *et al*. [11] (GEO GSE284197), focusing on ZNF219 and the concordant comparison targets ARX and NR2E1. Each target had three guides passing the source study’s knockdown filter, defined as target log_2_ fold change ≤ −0.4 in at least one cell class. We followed the analytical framework reported in Ding *et al*. [11] closely to reproduce their published results before applying PETRO.

For cell-state composition, we used day-7 cells from four donors and 3,137 non-targeting (NT) control cells. We analyzed four broad cell classes for all three targets and eight finer cell states for ZNF219. Counts for each class or state were summed by donor and perturbation label and analyzed using beta-binomial regression in DCATS v.1.4.0 [24] with donor as a covariate. Guide-level analyses compared cells carrying each individual guide with NT controls, whereas pooled analyses used the source study’s target-level perturbation labels. We report log_2_ odds ratios relative to NT, model-based 95% Wald confidence intervals and likelihood-ratio test *P* values.

For the STICR lineage analysis, we retained clones containing at least three cells from the eight donors with evaluable ZNF219 lineage data. For each donor and perturbation condition, we calculated the fraction of clones biased toward inhibitory neurons. ZNF219 g1, g3 and the pooled ZNF219 condition were compared with NT controls using exact two-sided Wilcoxon signed-rank tests paired by donor.

For transcriptional analyses, counts from inhibitory neurons carrying a single guide were summed into pseudobulks by donor and perturbation label. We fitted one PyDESeq2 v.0.4.12 model [25] per target at either the guide or pooled-target level, comparing each perturbation label with NT controls. For each outcome gene, PETRO combined the three guide-level log_2_ fold-change estimates and their covariance from the same DESeq2 model using pointwise parametric inference. The 63-gene ZNF219 comparison used the inhibitory-neuron network reported by the source study. Guide sign agreement was calculated relative to the pooled estimate, and genes were classified according to whether all three guides agreed in sign or one guide opposed the other two.

For gene set enrichment analysis, genes from each guide-level, pooled and PETRO profile were ranked by signed *z*-score and analyzed using fgsea v.1.32.4 [26]. We tested the MSigDB 2026.1.Hs Hallmark [27], Reactome [28] and Gene Ontology Biological Process [29] collections together with the five stress-response gene sets from the source study, retaining gene sets containing 10-500 genes. Significance was defined as Bonferroni-adjusted *P <* 0.10, with correction performed separately within each target, analysis and gene-set collection. To assess guide support for the pooled interpretation, we first identified terms significant in the pooled profile and then recorded whether the corresponding guide-level and PETRO profiles were significant in the same or opposite direction.

## Data availability

Raw sequencing data for the Southard RPE-1 and Hs27 datasets are available from the Sequence Read Archive under accession PRJNA1108254. Processed RPE-1 data are available in Zenodo records 15211972 and 15213619, and processed Hs27 data are available in Zenodo records 15213597 and 15200179. Raw HEK293FT data are available from the Gene Expression Omnibus under accession GSE269600, and processed data are available in Zenodo record 13238298. The K562 and A549 CRISPRi data and SCEPTRE results are available from Zenodo (doi:10.5281/zenodo.20722064 and doi:10.5281/zenodo.21136747). The A375 CRISPRa data are available from Figshare (doi:10.6084/m9.figshare.31119196). The cortical Perturb-seq and lineage data are available from the Gene Expression Omnibus under accession GSE284197.

## Code availability

The PETRO implementation and accompanying examples are available at https://github.com/zsun263/PETRO. Processed analysis tables will be released in a permanent repository before publication.

## Notes

### Competing Interest Statement

The authors have declared no competing interest.

## References

[1] Adamson, B. et al. A multiplexed single-cell CRISPR screening platform enables systematic dissection of the unfolded protein response. Cell 167, 1867–1882.e21 (2016).

[2] Xu, Z., Sziraki, A., Lee, J., Zhou, W. & Cao, J. Dissecting key regulators of transcriptome kinetics through scalable single-cell RNA profiling of pooled CRISPR screens. Nature Biotechnology 42, 1218–1223 (2024).

[3] Southard, K. M. et al. Comprehensive transcription factor perturbations recapitulate fibroblast transcriptional states. Nature Genetics 57, 2323–2334 (2025).

[4] Love, M. I., Huber, W. & Anders, S. Moderated estimation of fold change and dispersion for RNA-seq data with DESeq2. Genome Biology 15, 550 (2014).

[5] Barry, T., Mason, K., Roeder, K. & Katsevich, E. Robust differential expression testing for single-cell CRISPR screens at low multiplicity of infection. Genome Biology 25, 124 (2024).

[6] Jiang, L. et al. Systematic reconstruction of molecular pathway signatures using scalable single-cell perturbation screens. Nature Cell Biology 27, 505–517 (2025).

[7] Du, J.-H., Shen, M., Mathys, H. & Roeder, K. Uncovering causal relationships in single-cell omic studies with causarray. Briefings in Bioinformatics 27, bbag175 (2026).

[8] Rohatgi, N. et al. Seed sequences mediate off-target activity in the CRISPR-interference system. Cell Genomics 4, 100693 (2024).

[9] Drepanos, L. M. et al. Direct comparison of CRISPR knockout and interference with Perturb-seq. Preprint at 10.64898/2026.07.04.736492 (2026).

[10] Hartman, A. et al. Systematic identification of seed-driven off-target effects in Perturb-seq experiments. Preprint at 10.64898/2026.03.27.714658 (2026).

[11] Ding, J. W. et al. Dissecting gene regulatory networks governing human cortical cell fate. Nature 651, 732–742 (2026).

[12] Wu, P. & Zhou, X. Canon enables causal inference of downstream genes in single-cell CRISPR studies via instrumental variable analysis. Proceedings of the National Academy of Sciences 123, e2525359123 (2026).

[13] Han, C. Detecting invalid instruments using L1-GMM. Economics Letters 101, 285–287 (2008).

[14] Kang, H., Guo, Z., Liu, Z. & Small, D. Identification and inference with invalid instruments. Annual Review of Statistics and Its Application 12, 385–405 (2025).

[15] Christensen, O. P., Markham, A., Kang, H., Gabriel, E. & Pers, T. H. Causal effect estimation from trans-regulatory single-cell CRISPR screens. Cell Genomics 6, 101251 (2026).

[16] Beaton, A. E. & Tukey, J. W. The fitting of power series, meaning polynomials, illustrated on band-spectroscopic data. Technometrics 16, 147–185 (1974).

[17] Brown, B. C., Tokolyi, A., Morris, J. A., Lappalainen, T. & Knowles, D. A. Large-scale causal discovery using interventional data sheds light on gene network structure in K562 cells. Nature Communications 16, 9628 (2025).

[18] Sun, Z., Kang, H. & Keleş, S. Causal gene regulatory network inference from Perturb-seq via adaptive instrumental variable modeling. Preprint at 10.64898/2026.02.18.706642 (2026).

[19] Fox, J., Kleiber, C. & Zeileis, A. ivreg: instrumental-variables regression by 2SLS, 2SM, or 2SMM, with diagnostics. R package version 0. 6-8 https://CRAN.R-project.org/package=ivreg (2026).

[20] Benjamini, Y. & Hochberg, Y. Controlling the false discovery rate: a practical and powerful approach to multiple testing. Journal of the Royal Statistical Society: Series B (Methodological) 57, 289–300 (1995).

[21] Akana, R. V. et al. High-content CRISPR activation screens identify synthetically lethal RNA-based mechanisms to sensitize cancer cells to targeted T cell cytotoxicity. Nature Genetics 58, 841–853 (2026).

[22] Robinson, M. D., McCarthy, D. J. & Smyth, G. K. edgeR: a Bioconductor package for differential expression analysis of digital gene expression data. Bioinformatics 26, 139–140 (2010).

[23] Kowalski, M. H. et al. Multiplexed single-cell characterization of alternative polyadenylation regulators. Cell 187, 4408–4425.e23 (2024).

[24] Lin, X., Chau, C., Ma, K., Huang, Y. & Ho, J. W. K. DCATS: differential composition analysis for flexible single-cell experimental designs. Genome Biology 24, 151 (2023).

[25] Muzellec, B., Teleńczuk, M., Cabeli, V. & Andreux, M. PyDESeq2: a Python package for bulk RNA-seq differential expression analysis. Bioinformatics 39, btad547 (2023).

[26] Korotkevich, G. et al. Fast gene set enrichment analysis. Preprint at 10.1101/060012 (2021).

[27] Liberzon, A. et al. The Molecular Signatures Database hallmark gene set collection. Cell Systems 1, 417–425 (2015).

[28] Milacic, M. et al. The Reactome pathway knowledgebase 2024. Nucleic Acids Research 52, D672–D678 (2024).

[29] The Gene Ontology Consortium. The Gene Ontology knowledgebase in 2023. Genetics 224, iyad031 (2023).

