## Supplementary information for "PETRO enables robust estimation of on-target perturbation effects in Perturb-seq"

#### Supplementary Figures

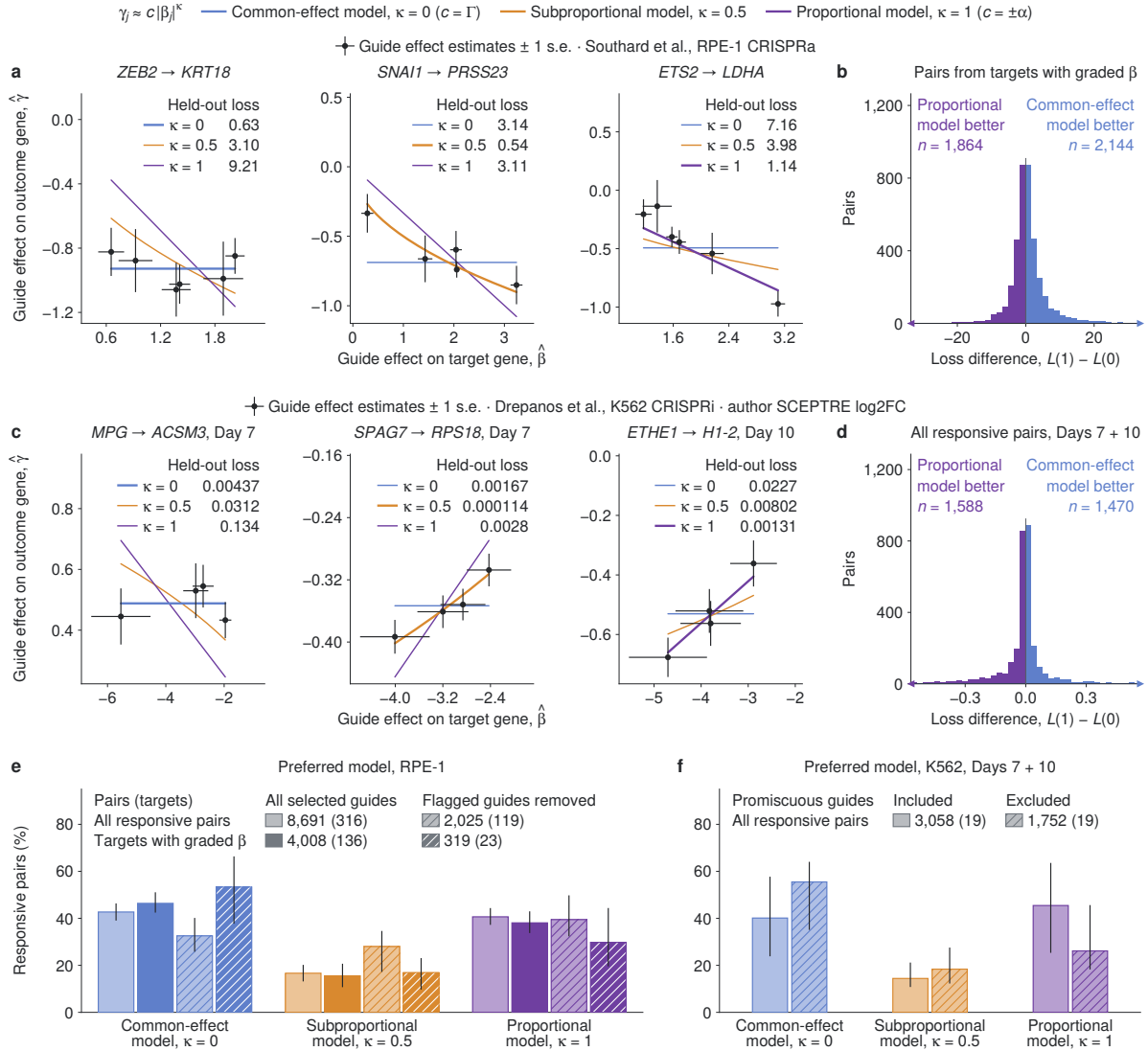

**Supplementary Figure S1. Held-out guide prediction across models for the on-target perturbation effect.** **a,c**, Illustrative RPE-1 CRISPRa and K562 CRISPRi target–outcome pairs, respectively. Points show estimated effects on target gene expression ($\hat{\beta}_j$) and guide effects on outcome expression ($\hat{\gamma}_j$), with $\pm 1$ standard error bars on both axes. RPE-1 estimates are raw ordinary least-squares coefficients; K562 estimates are author-reported SCEPTRE log<sub>2</sub> fold changes with delta-method standard errors. Common-effect ($\kappa = 0$, blue), subproportional ($\kappa = 0.5$, orange) and proportional ($\kappa = 1$, purple) curves are fitted to all displayed guides; the thicker curve marks the model with the lowest held-out loss. **b,d**, Loss differences $L(1) - L(0)$ for 4,008 responsive RPE-1 pairs from 136 targets with graded effects on target gene expression (**b**) and 3,058 K562 target–outcome–day pairs from 19 targets (**d**). Positive values favor the common-effect model; arrowheads mark pairs beyond the axis. Loss $L(\kappa)$ is the mean squared error of leave-one-guide-out predictions, with each squared error divided by the held-out estimate’s variance in RPE-1 and unweighted in K562. **e,f**, Preferred-model proportions, with pairs weighted equally. Light and dark fills denote all responsive RPE-1 pairs and the graded subset; hatching denotes removal of guides flagged by the source studies. K562 summaries combine days 7 and 10. Counts give pairs (targets). Error bars in **e,f** are 95% percentile intervals from 10,000 target bootstrap resamples. Guide selection, the definition of responsive pairs and the criteria for graded targets are described in Supplementary Note 2.

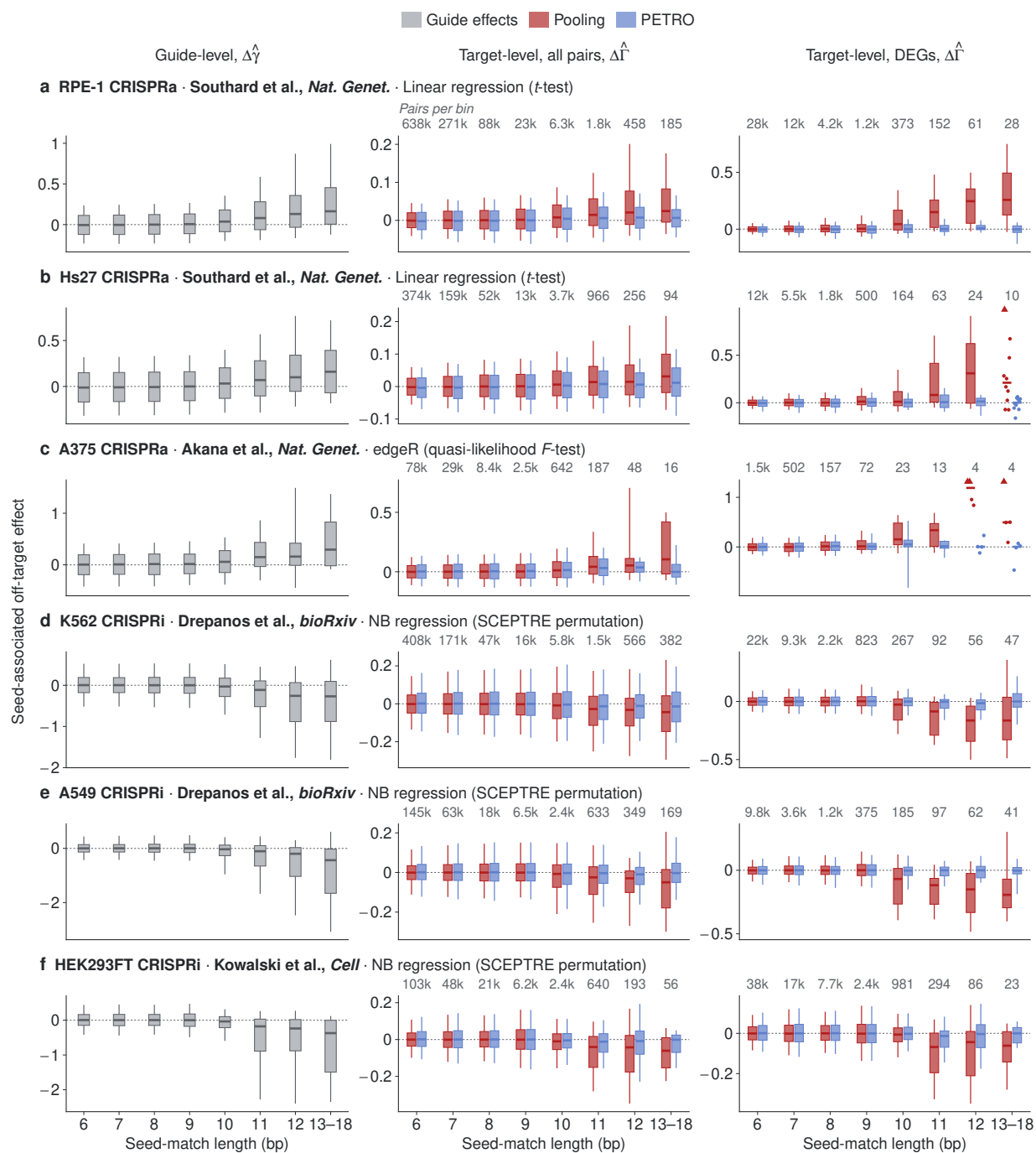

Supplementary Figure S2. See next page for caption.

**Supplementary Figure S2. PETRO reduces seed-associated bias across CRISPRa and CRISPRi screens and guide-level differential expression models.** **a,b**, RPE-1 and Hs27 CRISPRa [1]. **c**, A375 CRISPRa in monoculture [2]. **d,e**, K562 CRISPRi at days 4, 7, 10 and 14 combined, and A549 CRISPRi at day 7, respectively [3]. **f**, HEK293FT CRISPRi [4]. Row labels name the study, the guide-level differential expression model (NB, negative binomial) and, in parentheses, the test that calls differentially expressed genes (DEGs). Bias ($\Delta\hat{\gamma}$ for guides, $\Delta\hat{\Gamma}$ for targets) is the estimate minus a reference computed from the target’s guides without a $\geq 6$-bp PAM-proximal match to the outcome promoter (Methods), and at least one such guide is required. The reference is the pooled estimate of these guides for pooling and the inverse-covariance-weighted mean of their estimated effects for guide-level and PETRO estimates; the two references coincide under linear regression. Bins give the longest match of each guide (left) or among the target’s guides (middle and right). DEGs are outcomes with within-target Benjamini–Hochberg $q < 0.1$ [5] in the row’s test with the target’s guides pooled; the same DEGs are used for pooling and PETRO. Only guides that change target gene expression in the intended direction are used (**a,b,f**, Methods; **c**, $P < 0.05$ in the source study’s test of target gene expression; **d,e**, one-sided SCEPTRE $P < 0.05$); targets need at least five such guides and at least one guide retained by the published filter in **a,b**, and at least three elsewhere. Intended-target outcomes are omitted, as are pairs in which any guide has no expressing cells (**c–f**) and pairs in which guides with a $\geq 10$-bp match make up at least half of the target’s guides, for which the majority rule may not hold. Boxes span the 25th–75th and whiskers the 10th–90th percentiles, and lines mark medians. Bins with at most 10 pairs show individual pairs, and arrowheads mark pairs beyond the axis. Effects are $\log_2$ fold changes except in **a,b** (normalized expression units). Counts above bins give target–outcome pairs, summed over days in **d**.

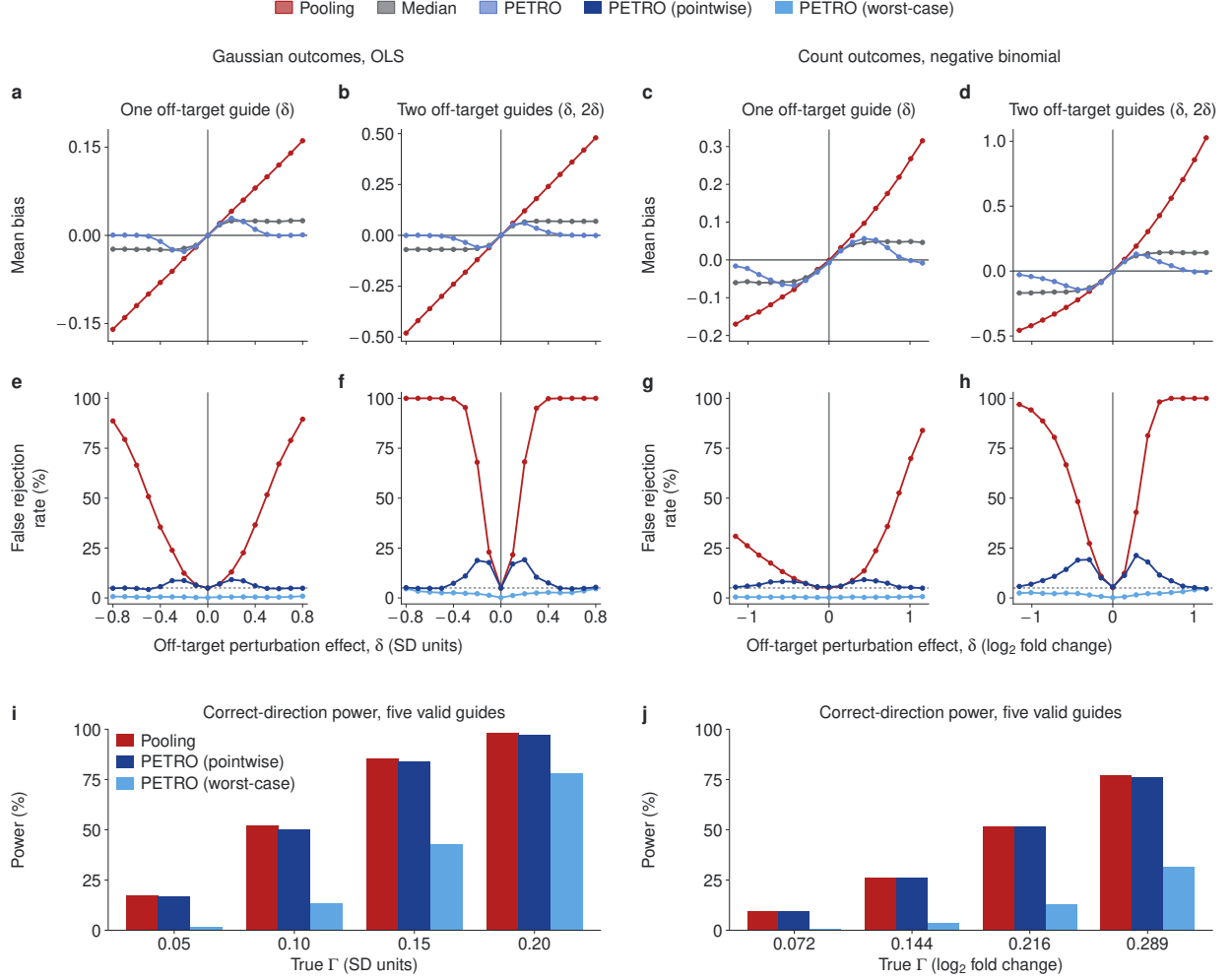

##### Supplementary Figure S3. Five-guide simulations with up to two off-target guides.

Each condition has five guides, 150 cells per guide, 900 shared non-targeting control cells and 10,000 replicates; the on-target perturbation effect is $\Gamma = 0$ in **a–h**. Off-target perturbation effects are $(0, 0, 0, 0, \delta)$ with one off-target guide and $(0, 0, 0, \delta, 2\delta)$ with two. Gaussian outcomes are fitted by ordinary least squares (OLS; **a, b, e, f, i**), with effects in residual standard deviation units; negative-binomial counts (**c, d, g, h, j**) have baseline mean 0.494 and size parameter 3.32 and are fitted by negative-binomial regression, with effects in $\log_2$ fold change. PETRO uses the adaptive cutoff and $q_{\max} = 0.4$. **a–d**, Mean bias of pooling, the guide median and PETRO. Pooling bias follows $\delta/5$ (**a**) and $3\delta/5$ (**b**) under OLS. Under the negative-binomial model, pooling averages counts, so its bias follows $\log_2\{(4 + 2^\delta)/5\}$ (**c**) and $\log_2\{(3 + 2^\delta + 2^{2\delta})/5\}$ (**d**), which are bounded below by $\log_2(4/5)$ and $\log_2(3/5)$ but grow without bound as $\delta$ increases. **e–h**, False rejection of $\Gamma = 0$ at two-sided $P \leq 0.05$; dotted lines mark 5%. Despite its smaller bias for negative $\delta$, negative-binomial pooling still rejects falsely (**g, h**). For Gaussian outcomes, pooling uses a Gaussian cell-level bootstrap and PETRO uses parametric inference with 9,999 null replicates. For negative-binomial counts, all tests use permutation inference with 999 permutations per replicate, and pooling uses the negative-binomial score test of SCEPTRE [6]. **i, j**, Correct-direction power with all five guides valid ($\Gamma = 0.05$–$0.20$ on the natural-log scale in **j**), defined as two-sided $P \leq 0.05$ and an estimate of the correct sign. Monte Carlo standard errors are at most 0.002 for bias and 0.5 percentage points for rates.

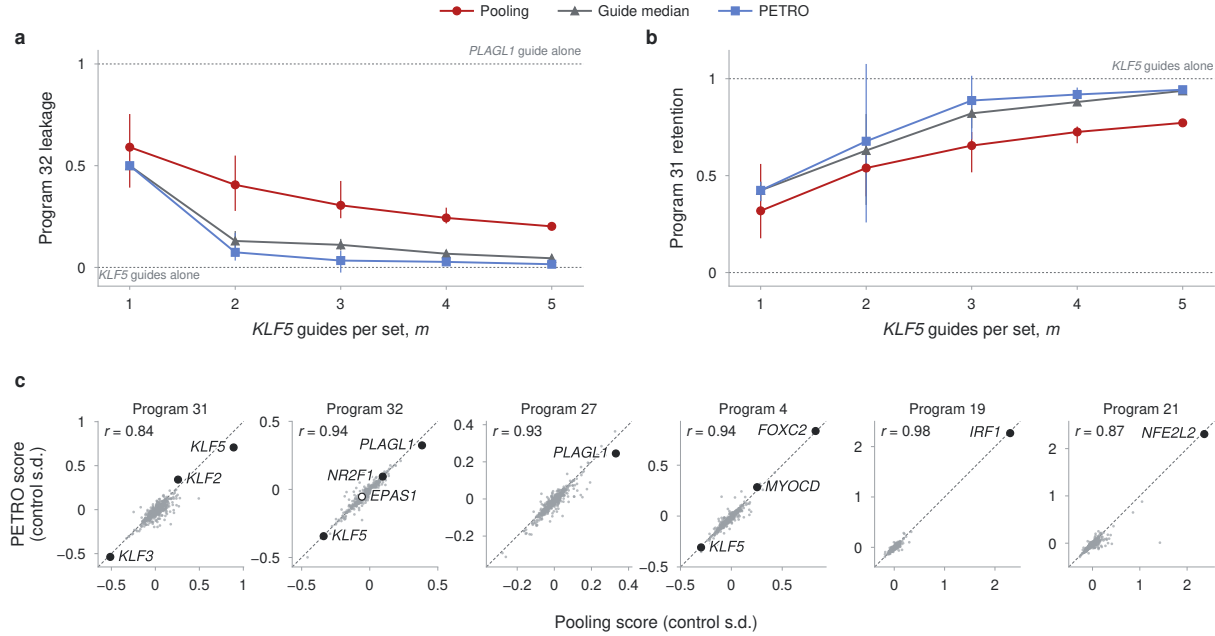

**Supplementary Figure S4. KLF5 benchmark and preservation of published target–program associations.** **a,b**, Semi-synthetic Hs27 KLF5 benchmark. One PLAGL1 guide was added to each of the 31 nonempty subsets of five KLF5 guides and treated as a KLF5 guide. Program 32 leakage (**a**) is the contaminated score minus the corresponding clean score, divided by the PLAGL1-only score minus that clean score. Program 31 retention (**b**) is the contaminated score divided by the corresponding clean score. Each method uses its own clean reference. Points show means and bars the minimum–maximum range across 5, 10, 10, 5 and 1 subsets with one through five KLF5 guides, respectively; dotted lines mark zero and one. **c**, Eligible target–program associations stated in the main text of Southard et al. [1]. Every candidate required at least three passing guides; 13 of 24 program-level statements were eligible. Atlas correlations alone did not qualify. Points show the signed loading-weighted scores of all 503 Hs27 targets under pooling and PETRO, in control-cell standard deviations (control s.d.); dashed lines mark identity, and $r$ is the Pearson correlation across targets. All eligible associations are displayed without a score or rank cutoff. Labelled filled points have the stated sign under both methods; the open point, EPAS1 activation of Program 32, has the opposite sign under both. EPAS1 activation was shown in the source study with a single guide that had no detectable activation of the target gene in Hs27 and did not pass guide filtering; neither method recovers it from the five remaining guides. In all panels, gene-level effects were aggregated before projection onto all published nonzero program loadings. Pooling uses uncapped full-covariance generalized least squares, which equals pooling cells across guides under linear regression; the guide median is unweighted; PETRO uses the fixed cutoff $c = 4.685$ and baseline weight cap $q_{\max} = 0.4$. With two guides, PETRO returns the median. Subset ranges and program scores are descriptive; no confidence intervals or $P$ values are shown. Benchmark definitions are given in the main Methods.

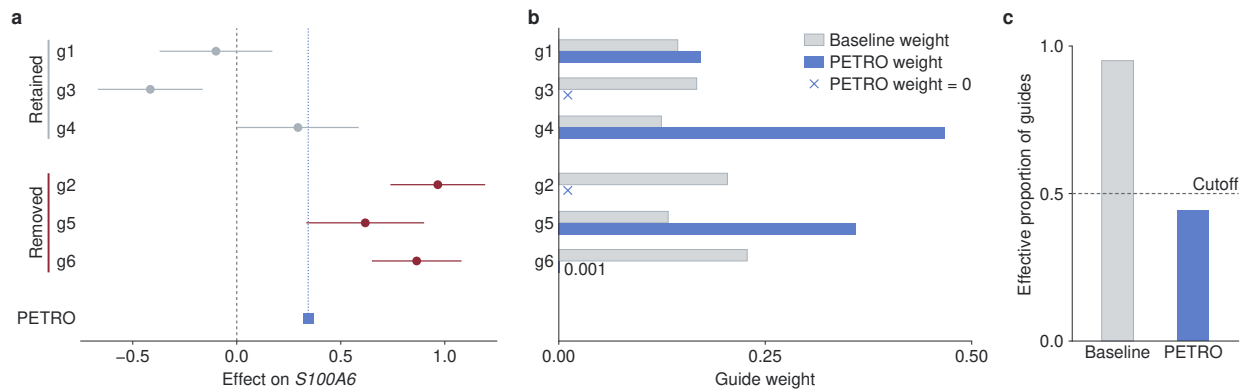

**Supplementary Figure S5. PETRO flags the MLXIP–S100A6 estimate for low precision-weighted agreement.** **a**, Estimated effects of six MLXIP guides on S100A6 in RPE-1 CRISPRa data [1]. Circles show raw ordinary least-squares coefficients, and horizontal bars show 95% normal confidence intervals. Grey and red indicate guides retained and removed by the published filter, respectively. The blue square shows the PETRO point estimate (0.344) from all six guides; the dashed line marks zero. The majority rule requires at least four valid guides. $F$ tests rejected the null hypothesis of equal guide effects in each of the 15 four-guide subsets (largest nominal $P = 0.0020$). **b**, Baseline weights $\hat{q}_j$ (grey) and final PETRO weights $\hat{w}_j$ (blue), with each set summing to one. Crosses mark zero final weights. **c**, Effective proportion of guides, calculated as $1/(6 \sum_j \hat{q}_j^2)$ for baseline weights and $1/(6 \sum_j \hat{w}_j^2)$ for final weights. Lower values indicate greater concentration of weight on fewer guides. The baseline and final proportions are 0.95 and 0.44, corresponding to 5.70 and 2.65 effective guides, respectively. The final proportion falls below the 0.5 cutoff (dashed line) for flagging low precision-weighted agreement.

### Supplementary Notes

#### Supplementary Note 1. PETRO estimation and inference

Here we describe how PETRO estimates and tests the on-target perturbation effect for each target–outcome pair $(A, Y)$. Let $j = 1, \dots, m$ index the guides designed to perturb $A$. Under the common-effect model, the guide effects are $\gamma_j = \Gamma + \delta_j$, where $\Gamma$ is the on-target perturbation effect and $\delta_j$ is the off-target perturbation effect of guide $j$. A guide is valid if $\delta_j = 0$, and the majority rule assumes that more than half of the guides are valid ($|\mathcal{V}| > m/2$, where $\mathcal{V} = \{j : \delta_j = 0\}$). For estimation, we first compute baseline weights from the precision of each guide and then downweight guides that disagree with the others using the Tukey bisquare loss. For inference, we provide two tests under the majority rule. The pointwise test uses the fitted bisquare weights to judge which candidate valid sets are plausible and generates null replicates from them. The worst-case test tests every candidate valid set of the smallest majority size and rejects the null hypothesis only if all are rejected. Either test can use parametric inference, which relies on the asymptotic normality of the estimates, or permutation inference, which shuffles the labels of the guides in a candidate valid set and the non-targeting controls and relies on their exchangeability.

##### 1.1 Guide summaries and baseline weights

The input consists of $m \geq 3$ estimated guide effects $\hat{\gamma} = (\hat{\gamma}_1, \dots, \hat{\gamma}_m)^\top$ from a guide-level differential expression (DE) model, such as linear or negative-binomial regression, and their estimated covariance $\hat{\Sigma}$. With two guides, PETRO returns their mean and performs no test. The equations and algorithms below use this covariance for both robust weighting and inference. For count models, robust weighting instead uses a working covariance evaluated at the guide median (Section 1.4).

Baseline weights account for differences in guide precision and dependence between estimates. We choose baseline guide weights using capped minimum-variance weighting:

$$\hat{\mathbf{q}} = \arg \min_{\mathbf{q}} \mathbf{q}^\top \hat{\Sigma} \mathbf{q}, \quad \sum_{j=1}^m q_j = 1, \quad 0 \leq q_j \leq q_{\max}. \quad (\text{S1})$$

Here $q_{\max} \in [1/m, 1]$ is the cap on each baseline weight. With no capping on $q_j$, this reduces to inverse-covariance weighting, and to inverse-variance weighting when $\hat{\Sigma}$ is diagonal.

##### 1.2 Robust estimation using the Tukey loss

We use the Tukey bisquare loss [7] to reduce the contribution of estimated guide effects that are distant from a candidate value $\theta$ of the on-target perturbation effect. For guide $j$, define $\hat{\tau}_j = \sqrt{\hat{\Sigma}_{jj}}$. The standardized residual and the bisquare loss are

$$u_j(\theta) = \frac{\hat{\gamma}_j - \theta}{\hat{\tau}_j}, \quad \rho_c(u) = \begin{cases} \frac{c^2}{6} \{1 - (1 - u^2/c^2)^3\}, & |u| < c, \\ c^2/6, & |u| \geq c. \end{cases} \quad (\text{S2})$$

Within each fit, the baseline weights $\hat{q}$ and residual scales $\hat{\tau}_j$ remain fixed as $\theta$ varies. The loss increases with the size of the standardized residual up to the cutoff $c$ and is constant beyond it, so a guide with $|u_j(\theta)| \geq c$ receives zero weight and is removed from the fit at that value of $\theta$. The objective used by PETRO is

$$\mathcal{L}(\theta) = \sum_{j=1}^m \hat{q}_j \hat{\tau}_j^2 \rho_c\{u_j(\theta)\}. \quad (\text{S3})$$

Differentiating the objective gives the bisquare weights $a_j(\theta)$ and the estimating score $F(\theta)$,

$$\begin{aligned} a_j(\theta) &= \{1 - u_j(\theta)^2/c^2\}^2 I\{|u_j(\theta)| < c\}, \\ F(\theta) &= \sum_{j=1}^m \hat{q}_j a_j(\theta) (\hat{\gamma}_j - \theta), \quad \mathcal{L}'(\theta) = -F(\theta), \end{aligned} \quad (\text{S4})$$

where $I\{\cdot\}$ is the indicator function. The bisquare weight is positive for $|u_j(\theta)| < c$ and decreases to zero as $|u_j(\theta)|$ approaches $c$. We use either the conventional cutoff $c = 4.685$, which gives 95% efficiency under normality when all guides are valid, or an adaptive cutoff $c_n = \{4.685^2 + \log \log(e^e + n^*)\}^{1/2}$, where $n^*$ is the effective number of cells of the least precise guide. The adaptive cutoff grows slowly with $n^*$, so that as the number of cells increases, valid guides are retained while invalid guides are still removed.

Because the bisquare loss is bounded, the objective is not convex and can have several local minima. We find candidate minima by solving $F(\theta) = 0$ and checking positive loss curvature:

$$\begin{aligned} F'(\theta) &= \sum_{j=1}^m \hat{q}_j \{1 - u_j(\theta)^2/c^2\} \{5u_j(\theta)^2/c^2 - 1\} I\{|u_j(\theta)| < c\}, \\ \mathcal{L}''(\theta) &= -F'(\theta). \end{aligned} \quad (\text{S5})$$

Between consecutive breakpoints $\hat{\gamma}_j \pm c\hat{\tau}_j$, $F$ is a polynomial of degree at most five, so we find its roots interval by interval with NumPy's polynomial root solver. We let $\mathcal{R}_0$ be the set of roots with $F'(\theta) < 0$. Among these candidate minima, we retain those supported by a strict majority of guides, that is, minima at which more than $m/2$ guides have positive bisquare weight:

$$\mathcal{R} = \left\{ \theta : F(\theta) = 0, F'(\theta) < 0, \sum_{j=1}^m I\{a_j(\theta) > 0\} > m/2 \right\}. \quad (\text{S6})$$

We then select the minimum in $\mathcal{R}$ closest to the unweighted guide median $\hat{\gamma}_{\text{med}} = \text{median}(\hat{\gamma}_1, \dots, \hat{\gamma}_m)$. If $\mathcal{R}$ is empty, or if two or more minima in $\mathcal{R}$ are equally close to the guide median, we return the guide median and flag the estimate as a median fallback.

At the selected minimum $\hat{\Gamma}_{\text{PETRO}}$, the final guide weights are the baseline weights multiplied by the bisquare weights and normalized:

$$\hat{w}_j = \frac{\hat{q}_j a_j(\hat{\Gamma}_{\text{PETRO}})}{\sum_{k=1}^m \hat{q}_k a_k(\hat{\Gamma}_{\text{PETRO}})}, \quad \hat{\Gamma}_{\text{PETRO}} = \sum_{j=1}^m \hat{w}_j \hat{\gamma}_j. \quad (\text{S7})$$

When PETRO returns the median, its guide-weight vector $\hat{\mathbf{w}}_{\text{med}}$ assigns weight 1 to the central ordered guide for odd $m$, or 1/2 to each of the two central guides for even $m$, and zero elsewhere; these weights are used to studentize the estimate in the tests below. Algorithm S1 summarizes the estimation procedure.

**Algorithm S1** PETRO estimation

**Input:** Estimated guide effects $\hat{\gamma} \in \mathbb{R}^m$, $m \geq 3$; positive-definite covariance $\hat{\Sigma}$; feasible baseline weight cap $q_{\max}$; cutoff $c > 0$

**Output:** Estimate $\hat{\Gamma}_{\text{PETRO}}$ and final guide weights $\hat{\mathbf{w}}$; median fallbacks are flagged

- 1: $\hat{\mathbf{q}} \leftarrow \arg \min_{\mathbf{q}} \mathbf{q}^\top \hat{\Sigma} \mathbf{q}$
  subject to $\sum_j q_j = 1$ and $0 \leq q_j \leq q_{\max}$
- 2: $\hat{\tau}_j \leftarrow \sqrt{\hat{\Sigma}_{jj}}$, $j = 1, \dots, m$
- 3: Find $\mathcal{R}_0 = \{\theta : F(\theta) = 0, F'(\theta) < 0\}$ ▷ Numerical root finding
- 4: **for** $\theta \in \mathcal{R}_0$ **do**
- 5: $n(\theta) \leftarrow \sum_j I\{a_j(\theta) > 0\}$
- 6: **end for**
- 7: $\mathcal{R} \leftarrow \{\theta \in \mathcal{R}_0 : n(\theta) > m/2\}$ ▷ Majority support
- 8: **if** $\mathcal{R} = \emptyset$ **then**
- 9: Flag the estimate as a median fallback
- 10: **return** $(\hat{\gamma}_{\text{med}}, \hat{\mathbf{w}}_{\text{med}})$
- 11: **end if**
- 12: $\hat{\Gamma}_{\text{PETRO}} \leftarrow \arg \min_{\theta \in \mathcal{R}} |\theta - \hat{\gamma}_{\text{med}}|$ ▷ Closest to the median
- 13: Compute normalized guide weights $\hat{\mathbf{w}}$ using Eq. (S7)
- 14: **return** $(\hat{\Gamma}_{\text{PETRO}}, \hat{\mathbf{w}})$

##### 1.3 Testing the on-target perturbation effect

**Null hypothesis.** We test whether the on-target perturbation effect is zero. Under the majority rule, fewer than half of the guides may be invalid, and their off-target perturbation effects are unrestricted. The null hypothesis is therefore composite:

$$H_0 : \quad \Gamma = 0 \quad \text{and} \quad \sum_{j=1}^m I\{\delta_j \neq 0\} < \frac{m}{2}.$$

With three guides, for example, $H_0$ includes $\delta = (0, 0, 0)^\top$ and every permutation of $(0, 0, d)^\top$, $d \in \mathbb{R}$.

It is convenient to write $H_0$ in terms of candidate valid sets. For a set $S$ of guides, let $H_{0,S}$ be the hypothesis that $\Gamma = 0$ and every guide in $S$ is valid, with the guides outside $S$ unrestricted and possibly also valid. Let $h = \lfloor m/2 \rfloor + 1$ be the smallest strict-majority size, and let $\mathcal{S}$ and $\mathcal{S}_h$ denote the families of candidate valid sets with $|S| \geq h$ and $|S| = h$, respectively. The null hypothesis $H_0$ holds exactly when $H_{0,S}$ holds for some $S \in \mathcal{S}$. Because every $S \in \mathcal{S}$ contains a set of size $h$, whose guides are then also valid, sets of size $h$ suffice to represent the same composite null:

$$H_0 = \bigcup_{S \in \mathcal{S}} H_{0,S} = \bigcup_{S \in \mathcal{S}_h} H_{0,S}. \tag{S8}$$

**Tests and types of inference.** With the adaptive cutoff, the pointwise test is asymptotically valid for each fixed set of off-target perturbation effects, and it can be more powerful. The worst-case test controls type I error without identifying the valid guides, provided each candidate valid set is tested validly, at a potential cost in power. Parametric inference relies on normal approximations, which can be inaccurate when cell numbers are small or expression is sparse [6]. Permutation inference avoids these approximations but requires exchangeability, as described below. We refer to each combination of test and type of inference by both names, for example the permutation worst-case test.

**Parametric and permutation inference.** Permutation inference shuffles the labels of the guides in $S$ and the non-targeting controls within prespecified strata, such as batch, keeping the number of cells with each label in each stratum. Outcomes, covariates and the labels of guides outside $S$ are unchanged, and the guide-level DE model is refitted to the permuted data exactly as to the observed data. The resulting null distribution is valid if, under $H_{0,S}$, the permuted labels are exchangeable within strata.

For the pointwise test, parametric inference redraws the estimates of the guides in $S$ from a normal distribution with mean zero and their estimated covariance. The estimates of the other guides keep their observed values, shifted only through their correlation with the redrawn estimates, for example through shared control cells, so their unknown effects never need to be estimated. For the worst-case test, parametric inference is a Wald test, described below.

**Pointwise test.** The pointwise test (Algorithm S2) compares the studentized PETRO estimate with its distribution under null replicates generated from a mixture of candidate valid sets. Each $S \in \mathcal{S}$ receives sampling weight $\omega(S)$ proportional to $\{\prod_{j \in S} \hat{a}_j \prod_{j \notin S} (1 - \hat{a}_j)\}^2$, where $\hat{a}_j$ is the bisquare weight of guide $j$ at $\hat{\Gamma}_{\text{PETRO}}$, so sets containing guides that the fit downweights receive little weight. The weights are normalized over all $S \in \mathcal{S}$ and held fixed across null replicates; if all unnormalized weights are zero, the test returns $P = 1$. For each null replicate, a set $S$ is sampled from $\omega$, null data (permutation) or null guide estimates (parametric) are generated for $S$, and PETRO is refitted with all guides. In the algorithm, PETRO denotes Algorithm S1, $\hat{\gamma}_S$ contains the estimated effects of guides in $S$, $\hat{\Sigma}_{SS}$ is their covariance block and $\hat{\Sigma}_{:,S}$ contains the corresponding columns of $\hat{\Sigma}$.

**Algorithm S2** PETRO pointwise test

**Input:** Estimated guide effects $\hat{\gamma}$ and covariance $\hat{\Sigma}$; baseline weight cap $q_{\max}$; cutoff $c$; number of null replicates $B \geq 1$; type $\in \{\text{permutation, parametric}\}$; for permutation, data $\mathcal{D}$, strata and guide-level DE model

**Output:** Two-sided $P$ value

```
1: $(\hat{\Gamma}_{\text{PETRO}}, \hat{\mathbf{w}}) \leftarrow \text{PETRO}(\hat{\gamma}, \hat{\Sigma}; q_{\max}, c)$
2: $T_{\text{obs}} \leftarrow \hat{\Gamma}_{\text{PETRO}} / \sqrt{\hat{\mathbf{w}}^\top \hat{\Sigma} \hat{\mathbf{w}}}$
3: $\hat{a}_j \leftarrow a_j(\hat{\Gamma}_{\text{PETRO}})$, $j = 1, \dots, m$
4: $r(S) \leftarrow \{\prod_{j \in S} \hat{a}_j \prod_{j \notin S} (1 - \hat{a}_j)\}^2$, $S \in \mathcal{S}$
5: if $\sum_{S \in \mathcal{S}} r(S) = 0$ then
6:   return 1
7: end if
8: $\omega(S) \leftarrow r(S) / \sum_{S' \in \mathcal{S}} r(S')$, $S \in \mathcal{S}$ ▷ Fixed across null replicates
9: for $b = 1, \dots, B$ do
10:   Sample $S \sim \omega(\cdot)$
11:   if type = permutation then
12:     Permute the labels of guides in $S$ and controls within strata
13:     Refit the guide-level DE model to the permuted data to obtain $(\tilde{\gamma}, \tilde{\Sigma})$
14:   else
15:     Draw $\mathbf{z}_S \sim N_{|S|}(\mathbf{0}, \hat{\Sigma}_{SS})$
16:     $\tilde{\gamma} \leftarrow \hat{\gamma} + \hat{\Sigma}_{:,S} \hat{\Sigma}_{SS}^{-1} (\mathbf{z}_S - \hat{\gamma}_S)$
17:     $\tilde{\Sigma} \leftarrow \hat{\Sigma}$ ▷ Covariance for fitting and studentization
18:   end if
19:   $(\tilde{\Gamma}_{\text{PETRO}}, \tilde{\mathbf{w}}) \leftarrow \text{PETRO}(\tilde{\gamma}, \tilde{\Sigma}; q_{\max}, c)$ ▷ Apply PETRO to all guides
20:   $T_b \leftarrow \tilde{\Gamma}_{\text{PETRO}} / \sqrt{\tilde{\mathbf{w}}^\top \tilde{\Sigma} \tilde{\mathbf{w}}}$
21: end for
22: $L \leftarrow \sum_{b=1}^B I\{T_b \leq T_{\text{obs}}\}$, $U \leftarrow \sum_{b=1}^B I\{T_b \geq T_{\text{obs}}\}$
23: return $\min\{1, 2[1 + \min(L, U)] / (B + 1)\}$
```

With the adaptive cutoff and consistent estimates of the guide effects, the bisquare weights approach one for valid guides and zero for invalid guides as the number of cells grows. Thus $\omega$ concentrates on the valid set, giving pointwise asymptotic validity for each fixed $\delta$ provided the parametric or permutation null distribution is valid.

**Worst-case test.** For each $S \in \mathcal{S}_h$, the worst-case test (Algorithm S3) refits the guide-level DE model with the guides in $S$ constrained to share one effect $\Gamma_S$, while guides outside $S$ keep their own coefficients, along with the covariates, offsets and controls. Under $H_{0,S}$, the shared effect is $\Gamma_S = 0$. Let $\hat{\Gamma}_S$ estimate this shared effect and $\hat{s}_S > 0$ be its sampling standard error from the chosen model and variance estimator. Each set is tested with the statistic $\hat{\Gamma}_S / \hat{s}_S$, either by permutation inference or, for parametric inference, by a Wald test, $p_S = 2\{1 - \Phi(|\hat{\Gamma}_S / \hat{s}_S|)\}$, with $\Phi$ the standard normal distribution function. The worst-case $P$ value is $P_{\text{worst}} = \max_{S \in \mathcal{S}_h} p_S$, so $H_0$ is rejected only if every $H_{0,S}$ is rejected.

**Algorithm S3** PETRO worst-case test

**Input:** Cell-level data $\mathcal{D}$; $m \geq 3$ guides targeting $A$; guide-level DE model; $\text{type} \in \{\text{permutation}, \text{parametric}\}$; for permutation, number of null replicates $B \geq 1$ and strata

**Output:** Two-sided worst-case $P$ value $P_{\text{worst}}$

```
1: $\mathcal{S}_h \leftarrow \{S \subseteq \{1, \dots, m\} : |S| = \lfloor m/2 \rfloor + 1\}$
2: for $S \in \mathcal{S}_h$ do
3:   Pool the guides in $S$ and fit the guide-level DE model to obtain $(\hat{\Gamma}_S, \hat{s}_S)$
4:   $T_{\text{obs},S} \leftarrow \hat{\Gamma}_S / \hat{s}_S$
5:   if $\text{type} = \text{permutation}$ then
6:     for $b = 1, \dots, B$ do
7:       Permute the labels of guides in $S$ and controls within strata
8:       Refit the pooled model to the permuted data to obtain $(\tilde{\Gamma}_S, \tilde{s}_S)$
9:       $T_{S,b} \leftarrow \tilde{\Gamma}_S / \tilde{s}_S$
10:    end for
11:    $L_S \leftarrow \sum_{b=1}^B I\{T_{S,b} \leq T_{\text{obs},S}\}$, $U_S \leftarrow \sum_{b=1}^B I\{T_{S,b} \geq T_{\text{obs},S}\}$
12:    $p_S \leftarrow \min\{1, 2[1 + \min(L_S, U_S)]/(B+1)\}$
13:  else
14:    $p_S \leftarrow 2\{1 - \Phi(|T_{\text{obs},S}|)\}$ ▷ Wald test
15:  end if
16: end for
17: $P_{\text{worst}} \leftarrow \max_{S \in \mathcal{S}_h} p_S$
18: return $P_{\text{worst}}$
```

Because $H_0$ is the union of the $H_{0,S}$, rejecting $H_0$ requires rejecting every $H_{0,S}$, which makes this an intersection–union test. Under $H_0$, at least one set $S_0 \in \mathcal{S}_h$ contains only valid guides, so $\Pr(P_{\text{worst}} \leq t) \leq \Pr(p_{S_0} \leq t)$ at every level $t$, and the right-hand side is at most $t$ when the test of $S_0$ is valid. Because $S_0$ is unknown, $P_{\text{worst}}$ controls type I error whenever the test of every candidate valid set is valid under its own null.

#### 1.4 Count models

In count models, the variance of a guide’s estimated effect depends on the mean expression in its cells and therefore on the guide’s own effect. To keep baseline weights and residual scales from depending on a guide’s off-target perturbation effect, for negative-binomial, edgeR [8] and DESeq2 [9] guide-level DE models, PETRO computes them from a working covariance $\hat{\Sigma}_w$, the inverse expected information of a fit in which every guide effect is fixed at the guide median and the remaining coefficients are re-estimated. The working covariance replaces $\hat{\Sigma}$ in Eq. (S1) and in $\hat{\tau}_j$ in Algorithms S1 and S2, whereas the null replicates and the studentized statistic use the sampling covariance $\hat{\Sigma}$ of the unrestricted fit. In parametric null replicates, both covariances are kept at their observed values. In permutation null replicates, both are recomputed from each refitted model.

#### 1.5 Proportional extension for the regulatory effect

The proportional extension of PETRO estimates the regulatory effect $\alpha$ by applying Algorithm S1 to the Wald ratios $\hat{\tau}_j = \hat{\gamma}_j / \hat{\beta}_j$ of the eligible guides, where $\hat{\beta}_j$ is the estimated effect of guide $j$ on target gene expression (Methods); the eligible guides are selected once, before testing. The sampling and working covariances of the Wald ratios are obtained by the delta method from the joint covariance of the estimated guide effects and effects on target gene expression. The sampling covariance is evaluated at the observed estimates. The working covariance, used for baseline weights and residual scales, replaces every Wald ratio in this formula by the median Wald ratio, so that, as in count models, baseline weights and residual scales do not depend on a guide’s own off-target perturbation effect.

Under $\alpha = 0$, valid guides have $\gamma_j = \alpha\beta_j = 0$, so the null hypothesis for a candidate valid set $S$ is again $\gamma_j = 0$ for $j \in S$, and $\alpha$ is tested with the same pointwise and worst-case tests. In the pointwise test, parametric null replicates redraw the guide effects in $S$ as in Algorithm S2, and the estimated effects on target gene expression keep their observed values, shifted only through their correlation with the redrawn effects; in permutation null replicates, only the guide labels in the guide-level DE model for the outcome are permuted, and the estimated effects on target gene expression are kept. The Wald ratios, their covariances and the baseline weights are then recomputed before PETRO is refitted. The worst-case test is the same as for $\Gamma$, applied to the eligible guides.

#### Supplementary Note 2. Comparison of models for the on-target perturbation effect

To ask whether guides that perturb their target more strongly also have larger effects on an outcome, we compared three models for these guide effects:

$$\gamma_j = \eta |\beta_j|^\kappa, \quad \kappa \in \{0, 0.5, 1\}, \quad (\text{S9})$$

where $\beta_j$ is the effect of guide $j$ on target gene expression and $\eta$ is a coefficient shared by all guides. The common-effect ($\kappa = 0$) and proportional ($\kappa = 1$) models correspond to those of the main text; the subproportional model ($\kappa = 0.5$) lies between them.

We used the RPE-1 CRISPRa guide effects of the main analysis (Methods) and the released SCEPTRE [6] log<sub>2</sub> fold changes for K562 CRISPRi at days 7 and 10 [3], keeping K562 guides that passed the authors’ quality control and reduced target gene expression (one-sided $P < 0.05$), in targets with at least four such guides. For each target–outcome pair, we held out each guide in turn, fitted $\eta$ to the remaining guides and predicted the held-out effect. A prediction was used only if the remaining guides showed an effect on the outcome at a six-standard-deviation threshold ($P \leq 1.97 \times 10^{-9}$; joint Wald test in RPE-1, Bonferroni-adjusted minimum SCEPTRE $P$ value in K562), and a pair was kept only if it had at least three such predictions. For each pair, the preferred model was the one with the smallest mean squared prediction error $L(\kappa)$; in RPE-1, fits used inverse-covariance weights and each squared error was divided by the held-out estimate’s variance. We repeated the comparison after removing guides flagged by the source studies (those removed by the published off-target filter in RPE-1 and promiscuous guides in K562) and, separately, in RPE-1 targets with graded effects on target gene expression, defined as effects that differed between guides (Benjamini–Hochberg $q \leq 0.05$) and spanned at least a twofold range, with no gap between adjacent effects wider than half the range and at least two guides inside its middle 60%.

No single model was preferred for all pairs. In every comparison, the common-effect and proportional models were each preferred for 26–55% of pairs (Supplementary Fig. S1e,f).

#### Supplementary References

- [1] Southard, K. M. *et al.* Comprehensive transcription factor perturbations recapitulate fibroblast transcriptional states. *Nature Genetics* **57**, 2323–2334 (2025).
- [2] Akana, R. V. *et al.* High-content CRISPR activation screens identify synthetically lethal RNA-based mechanisms to sensitize cancer cells to targeted T cell cytotoxicity. *Nature Genetics* **58**, 841–853 (2026).
- [3] Drepanos, L. M. *et al.* Direct comparison of CRISPR knockout and interference with Perturb-seq. Preprint at https://doi.org/10.64898/2026.07.04.736492 (2026).
- [4] Kowalski, M. H. *et al.* Multiplexed single-cell characterization of alternative polyadenylation regulators. *Cell* **187**, 4408–4425.e23 (2024).
- [5] Benjamini, Y. & Hochberg, Y. Controlling the false discovery rate: a practical and powerful approach to multiple testing. *Journal of the Royal Statistical Society: Series B (Methodological)* **57**, 289–300 (1995).
- [6] Barry, T., Mason, K., Roeder, K. & Katsevich, E. Robust differential expression testing for single-cell CRISPR screens at low multiplicity of infection. *Genome Biology* **25**, 124 (2024).
- [7] Beaton, A. E. & Tukey, J. W. The fitting of power series, meaning polynomials, illustrated on band-spectroscopic data. *Technometrics* **16**, 147–185 (1974).
- [8] Robinson, M. D., McCarthy, D. J. & Smyth, G. K. edgeR: a Bioconductor package for differential expression analysis of digital gene expression data. *Bioinformatics* **26**, 139–140 (2010).
- [9] Love, M. I., Huber, W. & Anders, S. Moderated estimation of fold change and dispersion for RNA-seq data with DESeq2. *Genome Biology* **15**, 550 (2014).
